# Exposure to *P. falciparum* and common cold viruses shape vaccine responses in early life

**DOI:** 10.64898/2026.07.06.736529

**Authors:** Florian Bach, George Sigal, Jacob Wohlstadter, Rayven Brown, Gaby Dunn, Teri Ngo, Brian Ngo, Cat Demos, Kenneth Musinguzi, Felistas Nankya, Abel Kakuru, Alyssa Sbarra, Grant Dorsey, Moses R Kamya, Saki Takahashi, Prasanna Jagannathan

## Abstract

Vaccine immunogenicity is consistently lower in low-income countries than in high-income settings, yet the factors driving this disparity remain incompletely understood. Using multiplexed electrochemiluminescence serology, we measured IgG and IgA responses to Expanded Program on Immunization (EPI) vaccines and common childhood viral infections in 89 Ugandan infants. We integrated detailed parasitological surveillance and maternal clinical data to examine how *P. falciparum* infection history, concurrent parasitemia, maternal gravidity, and early-life viral exposures shaped serological profiles. We found that infants mounted robust responses to most EPI vaccines, but critical gaps in protection persisted for diphtheria, measles and rubella. Children born to primigravid mothers had lower antibody levels at 8 weeks of age, independent of placental malaria and only partially explained by maternal age. Contrary to expectation, cumulative *P. falciparum* exposure was positively associated with antibody concentrations to diphtheria and varicella, and concurrent parasitemia was positively correlated with responses to multiple antigens. Seroconversion to rhinovirus C was associated with higher IgG and IgA levels to several vaccines. Together, these findings suggest that common microbial exposures during infancy, including respiratory viruses and *P. falciparum* may positively modulate vaccine responsiveness.

## Introduction

Many vaccines are less effective in low-income countries than high-income countries^1^. *Plasmodium spp.* exposure is one proposed contributor to this disparity, but the literature linking malaria parasites to heterologous vaccine responses is inconsistent and sparse^2^, with most vaccines supported by only a handful of studies. Across this limited evidence, *P. falciparum* infection at the time of vaccination is generally either unassociated with vaccine response or modestly negatively associated^3^. More recent data suggest that P*. falciparum* transmission intensity is associated with reduced durability of vaccine-induced immunity^4^, raising the possibility that ongoing parasite exposure after vaccination, not just infection at the time of vaccination, erodes immune memory. Whether and how post-vaccination transmission shapes long-term vaccine responses remains critically understudied.

*P. falciparum* is only one pathogen of many that have been found to modulate immune responses to other antigens. Epidemiological interactions between evolutionarily distant influenza A virus and rhinovirus, leading to asynchronous temporal peaks of transmission, are well-established^5^. These population-level data have been supported by mechanistic experiments that implicate type-I interferon responses as mediating cross-protection by inducing a non-specific antiviral state in human airway epithelial cultures^6^. Interactions between viruses and parasites have also been observed: infection with the intestinal hookworm *Heligmosomoides bakeri* protected mice from challenge with respiratory syncytial virus^7^, also by inducing an antiviral state mediated by type-I interferons. Antiparasitic immune responses to *P. falciparum* have been linked to altered responses to SARS-CoV2, possibly affecting clinical outcome^8^ Taken together, these data highlight that many common pathogens can influence immune responses to divergent antigens, even spurring interest in “universal vaccines” that induce non-specific antimicrobial states that could be broadly protective^9^.

Infants, in their first year of life, experience high rates of viral illnesses; they also receive the largest number of vaccines of any age group^10^. To better understand the interplay between vaccine responses and childhood infection histories, we studied immune responses in a cohort of Ugandan infants receiving vaccines under the Expanded Program on Immunization (EPI). We leveraged multiplexed serology measurements to measure antibody responses against a range of vaccine targets, and to detect seroconversion to a panel of common viral pathogens. We also closely tracked *P. falciparum* incidence through active case finding. These data allowed us to explore how *P. falciparum* and viral infection histories shape vaccine responses in a real-world setting.

## Results

Participants were selected from an ongoing malaria chemoprevention clinical trial in Busia, Eastern Uganda (MICDRoP, NCT04978272) evaluating the effect of intermittent preventive therapy during the first two years of life on protective immunity to malaria. In MICDRoP, children were randomized to receive placebo for two years, dihydroartemisinin piperaquire (DP) for two years, or DP for one year followed by placebo for one year. To investigate vaccine responsiveness and exposure to common childhood viruses, we utilized three novel multiplexed serology panels (Table 1) from MESO SCALE DISCOVERY® to detect virus-specific IgG and vaccine-specific IgG and IgA.

**Table 1.** Viral and bacterial target antigens for the serology panels used in the study. Respiratory Panel 5 includes two array elements (or “Spots”) – PIV Mix and Influenza Mix – that are mixtures of antigens from different viral serotypes. The components of these mixes are resolved into individual spots in Respiratory Panel 6. Abbreviations used in the table include EV (enterovirus), hMPV (human metapneumovirus), RV (rhinovirus), PIV (parainfluenza virus), RSV (respiratory syncytial virus) and CoV-2 (SARS Coronavirus 2).

| Spot | Target Antigens in MSD Serology Panels |  |  |
| --- | --- | --- | --- |
|  | Respiratory Panel 5 | Respiratory Panel 6 | Vaccine Panel 1 |
| 1 | EV-71 | Influenza A/H1 | S. Pneumo. Mix (Types 1, 4, 14) |
| 2 | Empty | Influenza A/H3 | Measles |
| 3 | hMPV | Influenza B | Polio |
| 4 | EV-D68 | Empty | Mumps |
| 5 | RV-C | Empty | Varicella |
| 6 | PIV Mix (Types 1 to 4) | PIV 1 | Rubella |
| 7 | RSV | PIV 2 | Diphtheria |
| 8 | CoV-2 | PIV 3 | Tetanus |
| 9 | Empty | PIV 4 | Rotavirus |
| 10 | Influenza Mix (H1, H3, B) | Empty | Pertussis |

We generated data from plasma samples from 89 children at 8, 24 and 52 weeks of age.

70/89 children were selected randomly, equally from the DP 1 year and placebo groups; an additional 19 children were added based on high incidence of *P. falciparum* parasitemia and/or malaria. Participants were 45% male, 40% received DP in the first year of life; 47.7% experienced malaria in first year of life, 83.1% were exposed to *P. falciparum* parasitemia at least once in the first year of life (Table 2.) Documented vaccine uptake in this population was generally very good: all vaccines courses were completed by >88% of children (Table 3).

**Table 2.**
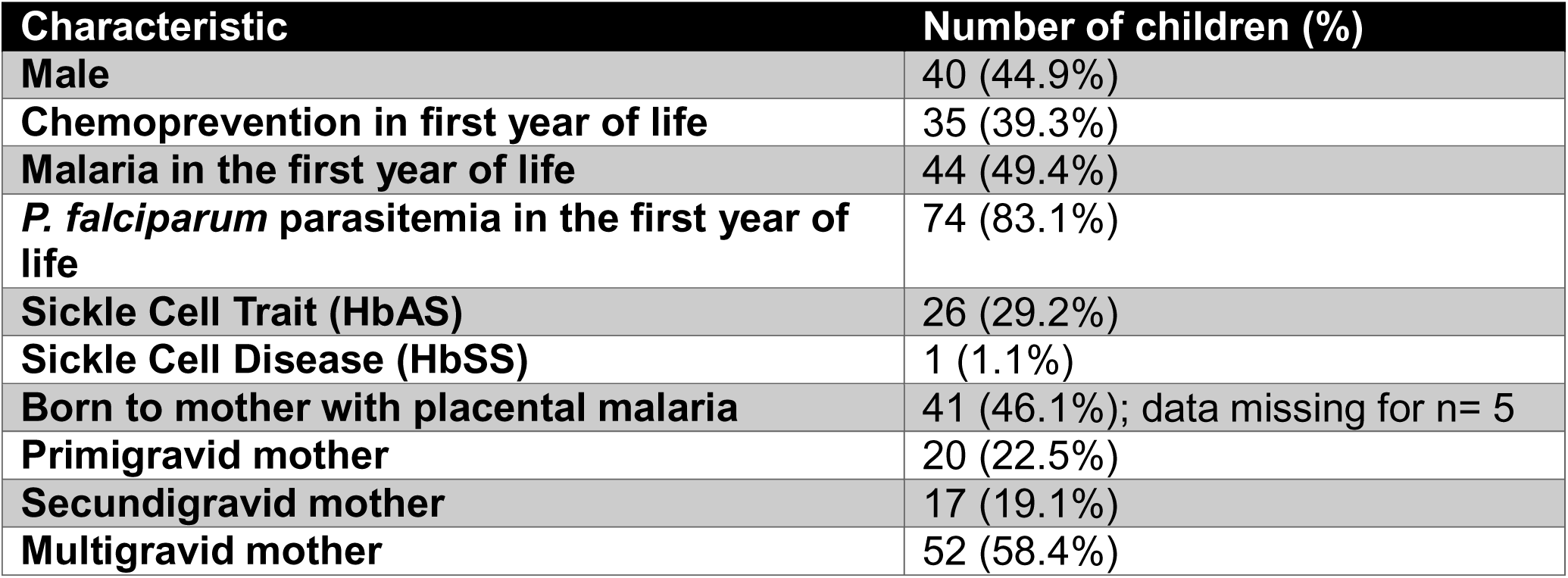
Demographic details of study cohort.

| Characteristic | Number of children (%) |
| --- | --- |
| Male | 40 (44.9%) |
| Chemoprevention in first year of life | 35 (39.3%) |
| Malaria in the first year of life | 44 (49.4%) |
| *P. falciparum* parasitemia in the first year of life | 74 (83.1%) |
| Sickle Cell Trait (HbAS) | 26 (29.2%) |
| Sickle Cell Disease (HbSS) | 1 (1.1%) |
| Born to mother with placental malaria | 41 (46.1%); data missing for n= 5 |
| Primigravid mother | 20 (22.5%) |
| Secundigravid mother | 17 (19.1%) |
| Multigravid mother | 52 (58.4%) |

**Table 3.** Documented vaccine uptake in the study cohort, by vaccine card and parental recall.

|  | complete | partial | none |
| --- | --- | --- | --- |
| **DTaP-Hib** | 81(91%) | 4(4.5%) | 4(4.5%) |
| **measles rubella** | 79(88.8%) | 0(0%) | 10(11.2%) |
| **Oral polio** | 81(91%) | 4(4.5%) | 4(4.5%) |
| **Oral polio at birth** | 84(94.4%) | 0(0%) | 5(5.6%) |
| **PCV10** | 82(92.1%) | 3(3.4%) | 4(4.5%) |
| **Rotarix** | 82(92.1%) | 2(2.2%) | 5(5.6%) |
| **Sabin polio** | 83(93.3%) | 0(0%) | 6(6.7%) |

### Infants mount robust vaccine responses to most EPI vaccines but a significant number of children remain below protective threshold at 52 weeks

First, we compared antibody concentration through time for each vaccine. For vaccines given at 6, 10 and 14 weeks (DTaP-Hib, PCV10, Rotarix, Oral polio) IgG to most antigens increased significantly from 8 weeks to 24 and 52 weeks. There were two exceptions, namely rotavirus and pertussis. For these antigens, immunization did not boost IgG responses relative to the maternal IgG antibody levels present at 8 weeks, however, IgA responses to these vaccines successively increased at 24 and 52 weeks relative to the lower maternal IgA background at 8 weeks. The vaccines given at 36 weeks (measles and rubella) showed a loss of maternal IgG between 8 and 24 weeks, and a strong increase in IgG between 24 and 52 weeks.

We next sought to determine whether childhood immunization induced protective antibody levels in this population. While protective thresholds have not been established for every vaccine antigen in our panel, we were able to estimate protective IgG thresholds for the following vaccines antigens: diphtheria, measles, Pneumococcus, polio, rubella and tetanus. These have established correlates of protection^11,12^, and available WHO serum standards for converting our assay results to accepted international concentration units. Of these vaccines, those given prior to 24 weeks (diphtheria, pneumococcus, polio and tetanus) produced protective levels of IgG in most children at 24 weeks. Only 23.6% of children retained levels of IgG against diphtheria toxoid ≥ 0.1IU / mL at 52 weeks of age (Fig 2B). Of note, 86.5% of children reached antibody levels ≥ 0.01 IU / mL which is thought to provide some degree of protection^13,14^. In contrast, IgG levels to tetanus, a component of the same combination vaccine that includes diphtheria, and polio remained at protective levels in greater than 90% of children at 52 weeks. The proportion of infants with protective levels of measles and rubella IgG at 52 weeks was more modest, 74.2% and 76.4% respectively. Postlicensure data from the United States Center for Disease Control and Prevention showed that a single dose of the MMR was 93% effective against measles and 97% effective against rubella^15^. It is unlikely that these levels of protection are reached in this cohort, given that only ∼75% reached protective levels, despite the fact that 88.8% of children received at least one dose.

**Figure 1:**
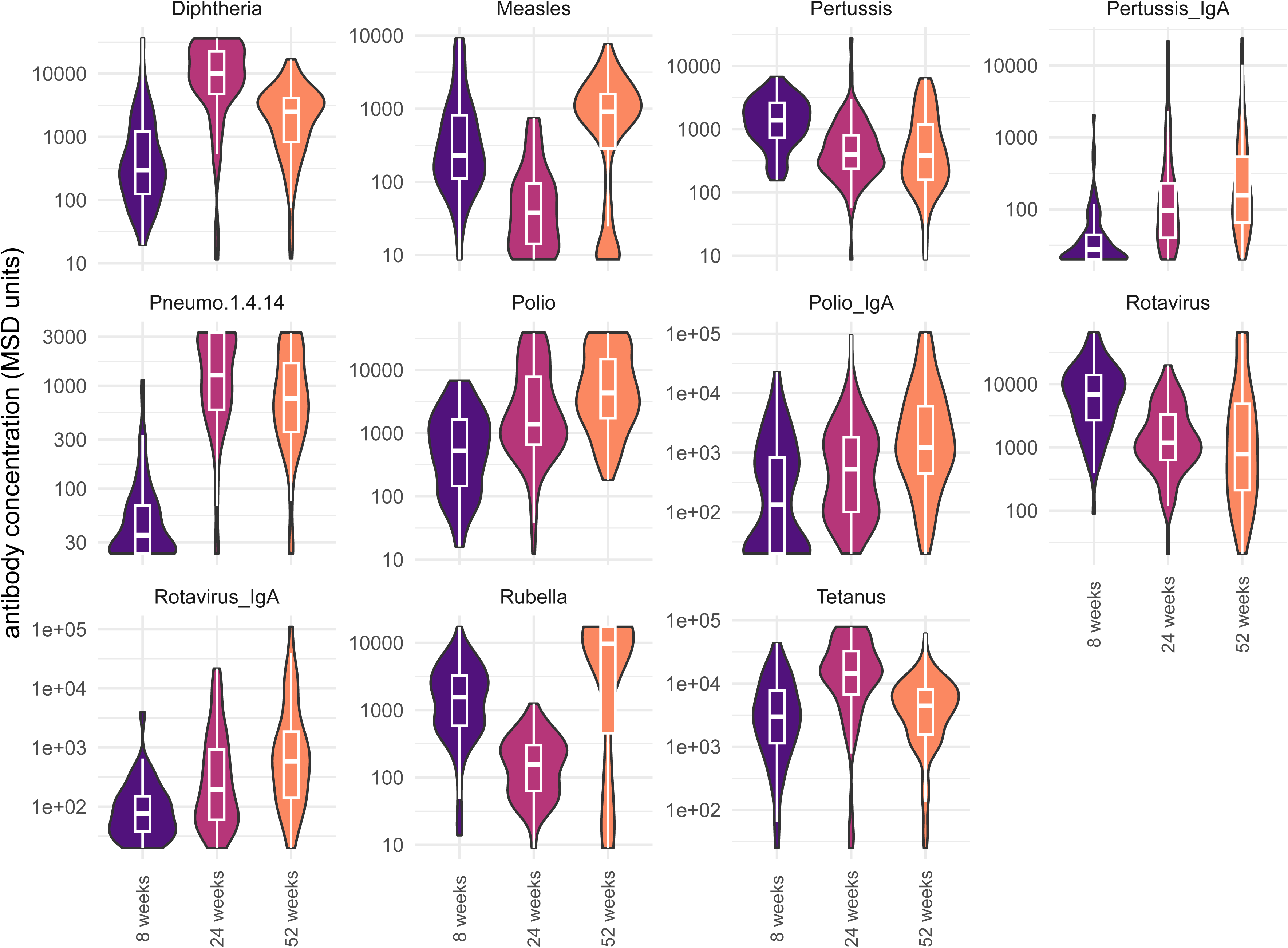
Longitudinal antibody responses to EPI vaccine antigens across the first year of life. Violin plots show plasma IgG (and IgA, where relevant) concentrations to vaccine antigens at 8, 24, and 52 weeks of age (n = 89). Boxes indicate median and interquartile range. Antibody concentrations are shown in MSD units on a log10 scale.

**Figure 2.**
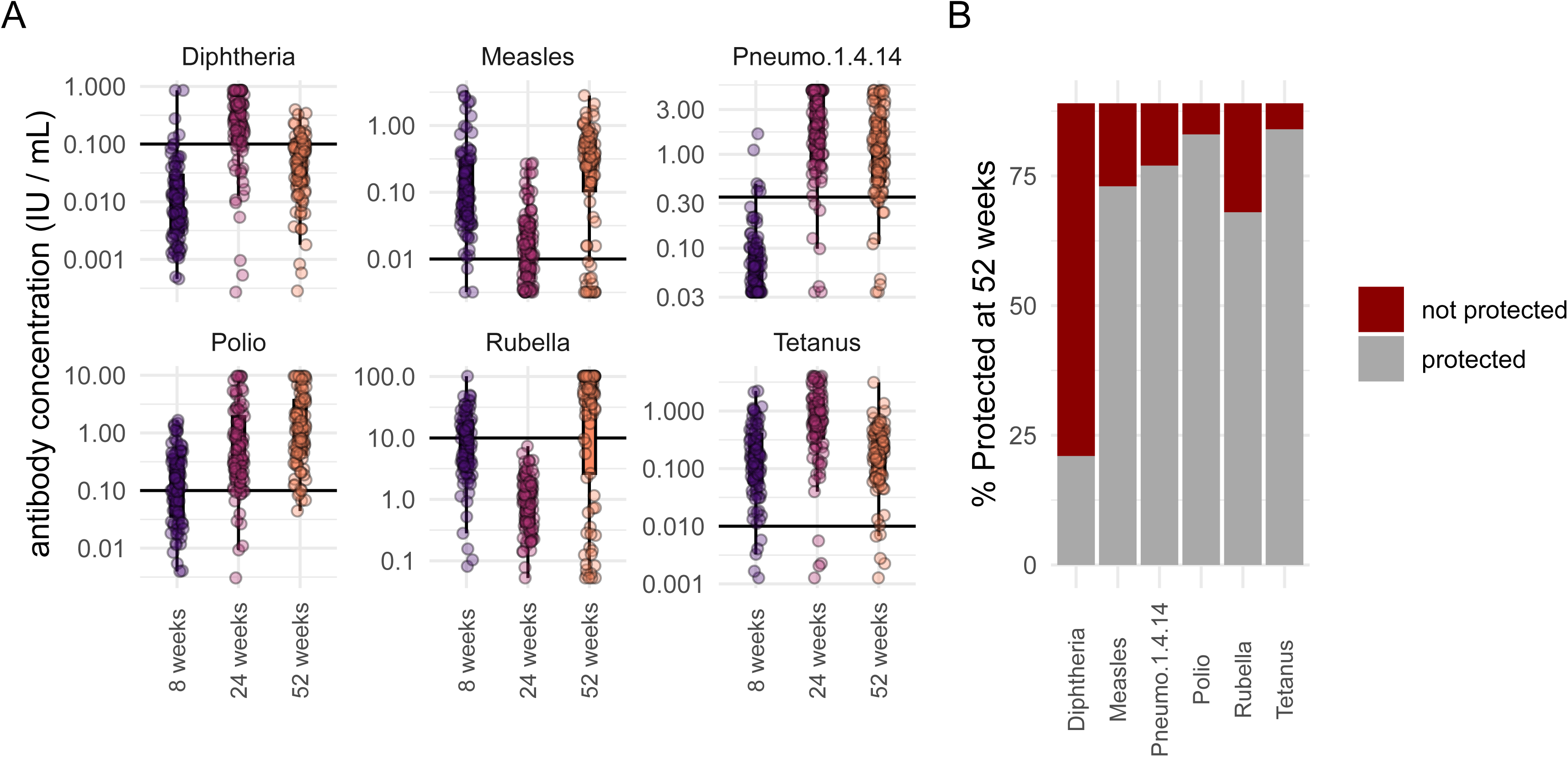
Vaccine antibody titers relative to established protective thresholds at 52 weeks of age. (A) Antibody concentrations (IU/mL) for diphtheria, measles, mix of pneumococcal serotypes 1.4.14, polio, rubella, and tetanus at 8, 24, and 52 weeks of age. Horizontal lines indicate established protective thresholds for each antigen (see Supplementary Table 3). Individual points represent single infants. (B) Percentage of infants reaching the protective threshold for each antigen at 52 weeks of age (grey, protected; red, not protected).

IgG levels to Pertussis declined progressively from 8 to 24 and 52 weeks. This suggests that infant immune responses fail to outpace the decline of maternal antibodies present at 8 weeks of age. In contrast, IgA responses to pertussis increased from 8 to 24 and 52 weeks of age. Interestingly, there was no significant correlation between pertussis IgG and pertussis IgA at any timepoint, while there were significant positive associations between IgG and IgA recognizing tetanus and diphtheria, other components of the same vaccine (DTPHib; Fig. S1).

Next, we tested associations between antibody concentrations at 8 weeks and later timepoints. Associations between 8 and 24 week concentrations were mixed, ranging from non-significant (PCV10 IgG, polio IgG, Rotavirus IgA, tetanus) to modest (Rotavirus IgG, polio IgA *ρ* ≥ 0.35) to very strong (measles, rubella *ρ* ≥ 0.8), shown in figure S2A. Interestingly, comparing antibody concentrations between 24 and 52 weeks showed no significant associations at all (Fig. S2B). Likely, these data reflect the waning of maternal antibodies, still strongly present at 8 and 24 weeks, and thus contributing to correlation between these timepoints. In addition, this suggests that durability of antibody responses cannot be predicted from peak responses shortly after vaccination in infants.

Collectively, these results suggest that infants in our Ugandan study cohort mount robust antibody responses to the majority of vaccine antigens, though notably IgG responses to diphtheria were not durable, leaving more than 75% of children susceptible at one year of age.

### Multiplexed Serology reveals high incidence of common childhood viral infections

Next, we measured seroconversion to common childhood viral pathogens, utilizing additional MSD arrays. We found evidence of seroconversion against a wide variety of viruses (Figure 3A and S3). The highest cumulative seroincidence (Fig 3B) was seen for rhinovirus C (61.8%) followed by enterovirus 71 (44.9%) and SARS CoV2 and respiratory syncytial virus (RSV) (both at 43.8%). These results are consistent with a high frequency of clinically reported upper respiratory tract infections (n=397) and diarrheal illness (n=153) in the first year of life in these 89 infants. There were 10 instances of mumps seroconversion in this cohort, though, interestingly, there were no clinical cases of mumps reported. Indeed, there was only one documented case of mumps in the parent clinical trial of 924 children, suggesting high rates of asymptomatic or undiagnosed mumps infections in infancy and early childhood.

**Figure 3.**
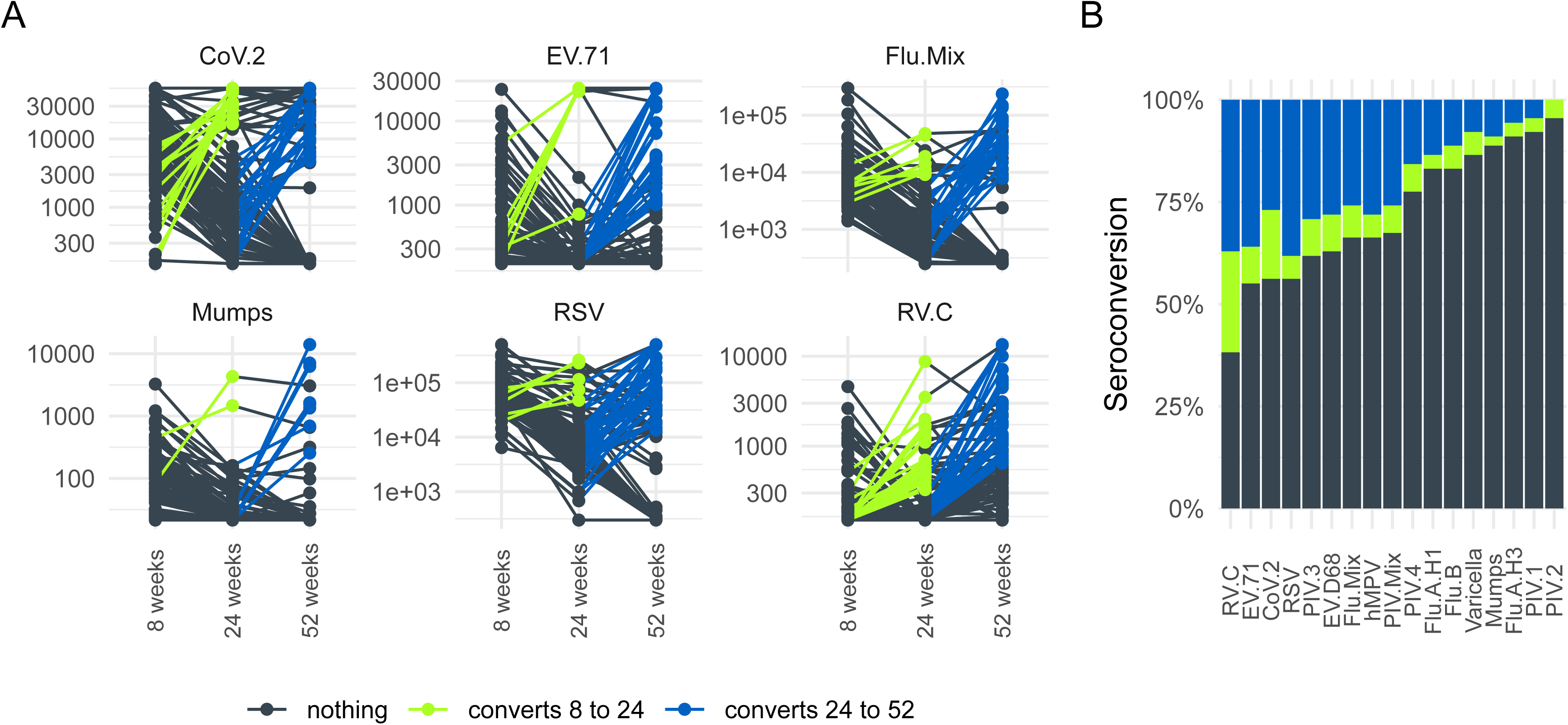
Cumulative seroconversion to common childhood viruses by one year of age. (A) Individual antibody trajectories from 8 to 24 and 52 weeks of age for select antigens: SARS-CoV-2 (CoV.2), enterovirus 71 (EV.71), influenza A and B mix (Flu.Mix), mumps, respiratory syncytial virus (RSV), and rhinovirus C (RV.C). Lines are colored by seroconversion status: dark grey, no seroconversion; green, seroconversion between 8 and 24 weeks; blue, seroconversion between 24 and 52 weeks. (B) Cumulative seroconversion by 52 weeks of age across all viral antigens assessed, ordered by overall seroconversion rate.

### Gravidity, but not placental malaria, is associated with lower antibody levels at 8 weeks

We used multivariate regression to test associations between antibody concentrations across all timepoints and various demographic factors. We found no associations between antibody titers and sex assigned at birth, maternal education level or household wealth category. The mothers of the children participating in this study were part of a clinical trial that tested alternatives to the standard of care for malaria prevention in pregnancy, intermittent preventative treatment (IPTp). Compared to the standard of care (sulfadoxine pyrimethamine, SP) maternal IPTp with dihydroartemisinin (DP) or a combination of DP+SP was not associated with differences in antibody titers at any timepoint.

Interestingly, we found that children born to primigravid mothers had lower levels of antibodies to several viral and vaccine antigens at 8 weeks (Fig. 4). After multiple testing correction, levels of IgG recognizing measles, rotavirus, tetanus, varicella and parainfluenza virus 2 (PIV2) were all significantly lower in infants born to primigravid mothers compared to *secundi*- and/or *multigravidae*. Surprisingly, the effect of gravidity appeared unrelated to placental malaria, as all interactions remained statistically significant after adjusting for the presence of histopathological findings. This finding was only partially associated with maternal age: adjusting for maternal age, we still found significant effects of gravidity on antibody levels to PIV2 and tetanus.

**Figure 4.**
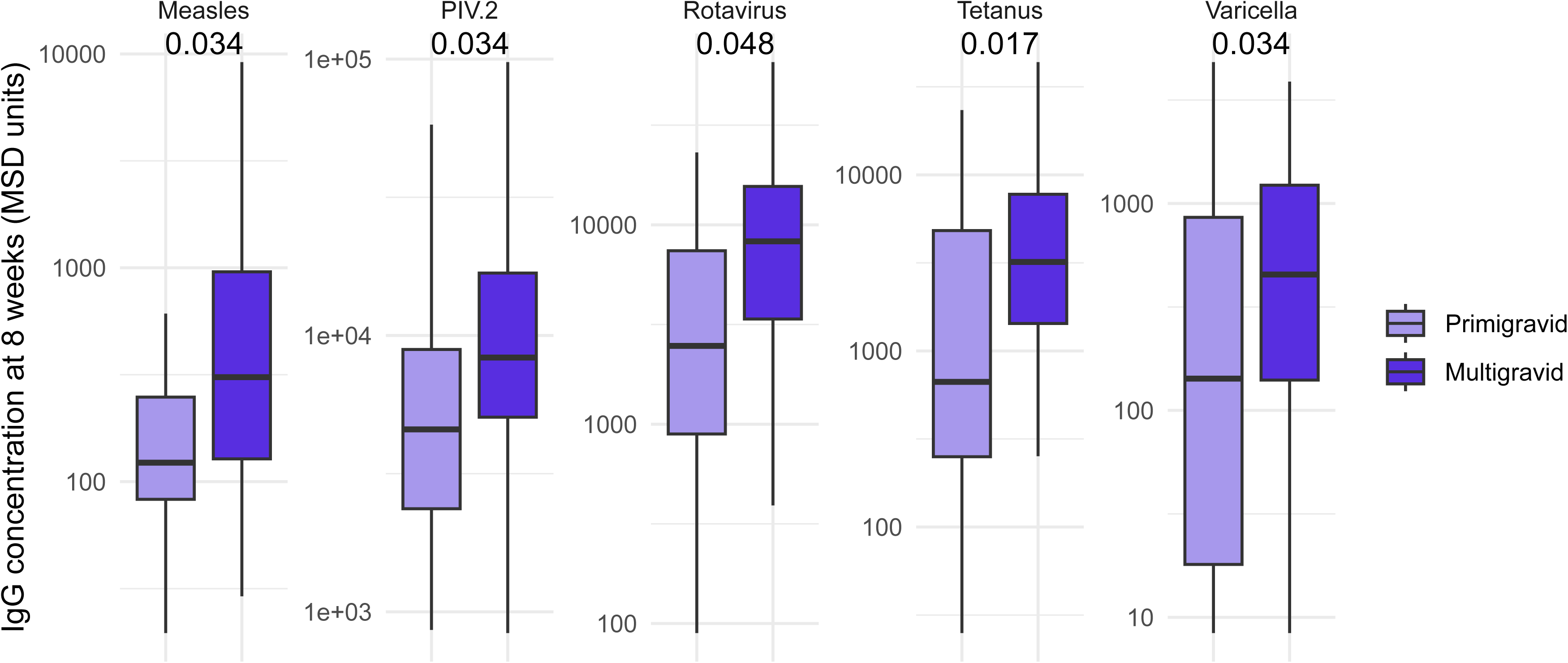
Maternal gravidity is associated with infant antibody concentrations at 8 weeks of age. IgG concentrations to measles, parainfluenza virus 2 (PIV.2), rotavirus, tetanus, and varicella at 8 weeks of age, stratified by maternal gravidity (primigravid, secundigravid, multigravid). All antibodies with adjusted p values < 0.1 are shown. Boxes show median and interquartile range.

In summary, we found gravidity was the only maternal demographic factor that was significantly associated with antibody concentrations in infants. This affect was only partially mediated by maternal age and was independent of IPTp and placental malaria.

### *P. falciparum* parasitemia at time of measurement, but not infection history, is associated with antibody concentrations to viral and vaccine antigens

We next sought to test whether *P. falciparum* infection history was associated with either viral or vaccine antibodies. To do this, we leveraged the frequent active case finding which was carried out during the clinical trial. Participants were scheduled to attend clinic visits every 28 days to receive study drug or placebo, at which point blood smear and/or qPCR was performed to determine *P. falciparum* parasite density. This allowed us to categorize each month for each child into one of three categories: uninfected, asymptomatic, or symptomatic parasitemia. We counted the number of parasitemic months and/or symptomatic malaria episodes to summarize infection history in various time windows for each child. We then used multivariate regression to test the relationship between the incidence of parasitemia or malaria in a given time window on antibody concentration. We adjusted for parasitemia at time of measurement, because we found consistent, albeit modest, positive correlations between parasite density and antibody concentrations generally, including both viral and vaccine antigens (see figures 5 and S4). To test the impact of infection history independent of parasitemia, we ran separate models for serological data at 24 weeks, testing infection histories up to 14 weeks, the timepoint for the final dose of most EPI vaccines, as well as complete infection histories up to 24 weeks. We also tested the association between infection histories up to 52 weeks on antibody concentrations at 52 weeks.

**Figure 5.**
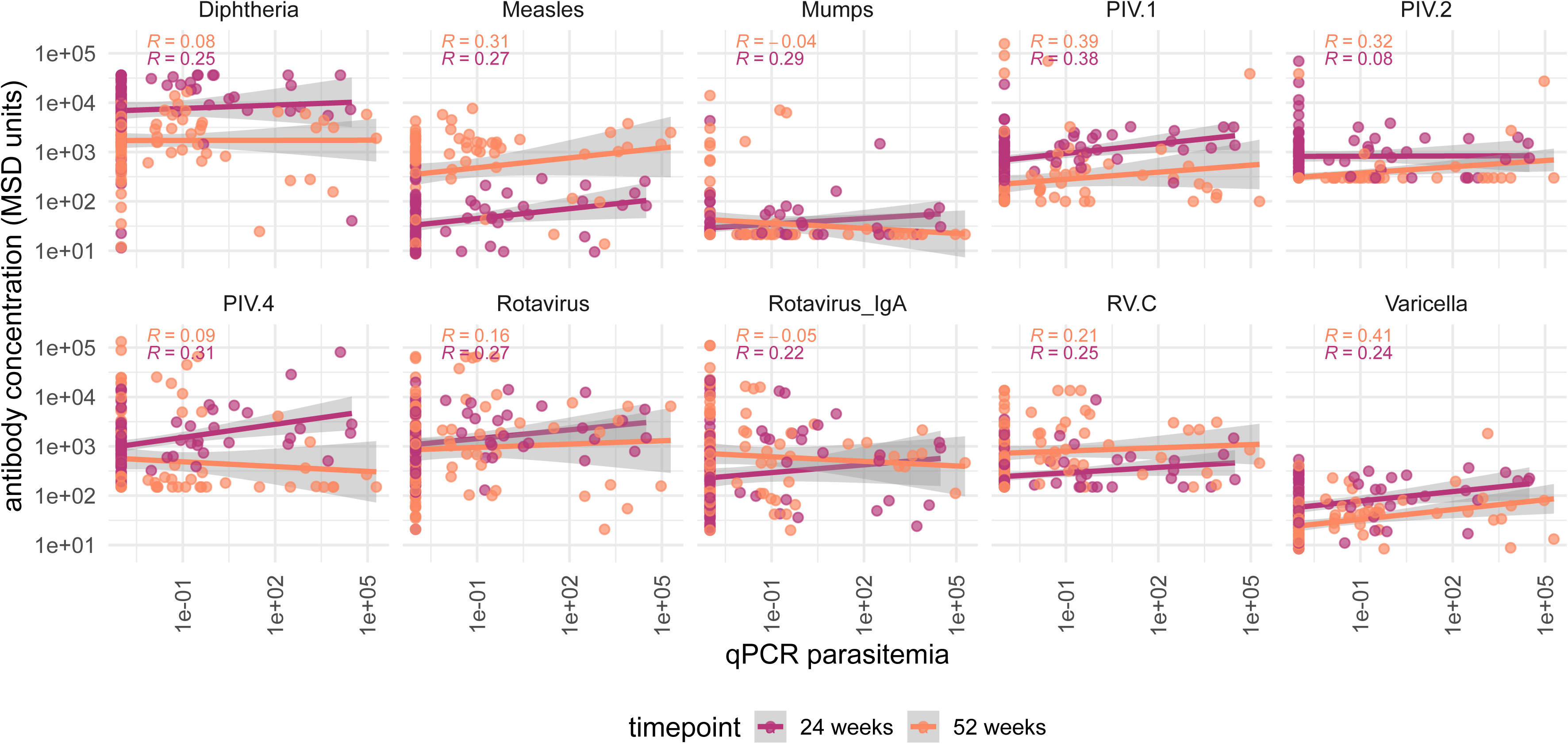
Antibody concentrations to multiple vaccine and viral antigens correlate with concurrent *P. falciparum* parasitemia. Antibody concentrations (MSD units, log10 scale) plotted against qPCR-quantified *P. falciparum* parasitemia at 24 weeks (magenta) and 52 weeks (orange) of age, for diphtheria, measles, mumps, PIV.1, PIV.2, PIV.4, rotavirus IgG and IgA, rhinovirus C (RV.C), and varicella. Lines show linear regression fit with 95% confidence intervals; R values indicate Spearman correlation coefficients for each timepoint.

Parasite prevalence or symptomatic malaria incidence before 14 weeks was not significantly associated with antibody titers at 24 or 52 weeks. We found that the number of parasitemic months in the first year of life was positively associated with varicella IgG, diphtheria IgG and diphtheria IgA (Fig. 6A). Symptomatic malaria was only significantly associated with varicella IgG, where the number of parasitemic months in the first year was also associated with antibody levels. Based on our previous work^16^, we investigated the relationship between tetanus antibodies between 24 and 52 weeks and the number of parasitemic months and/or symptomatic malaria in the first year of life. Fort tetanus, we did not observe significant associations between antibody concentrations and the number of parasitemic months in the first 12 months of life (Fig. S5). It remains possible that frequent infections after infancy negatively impact the durability of tetanus antibody responses, as previously shown^4^.

**Figure 6.**
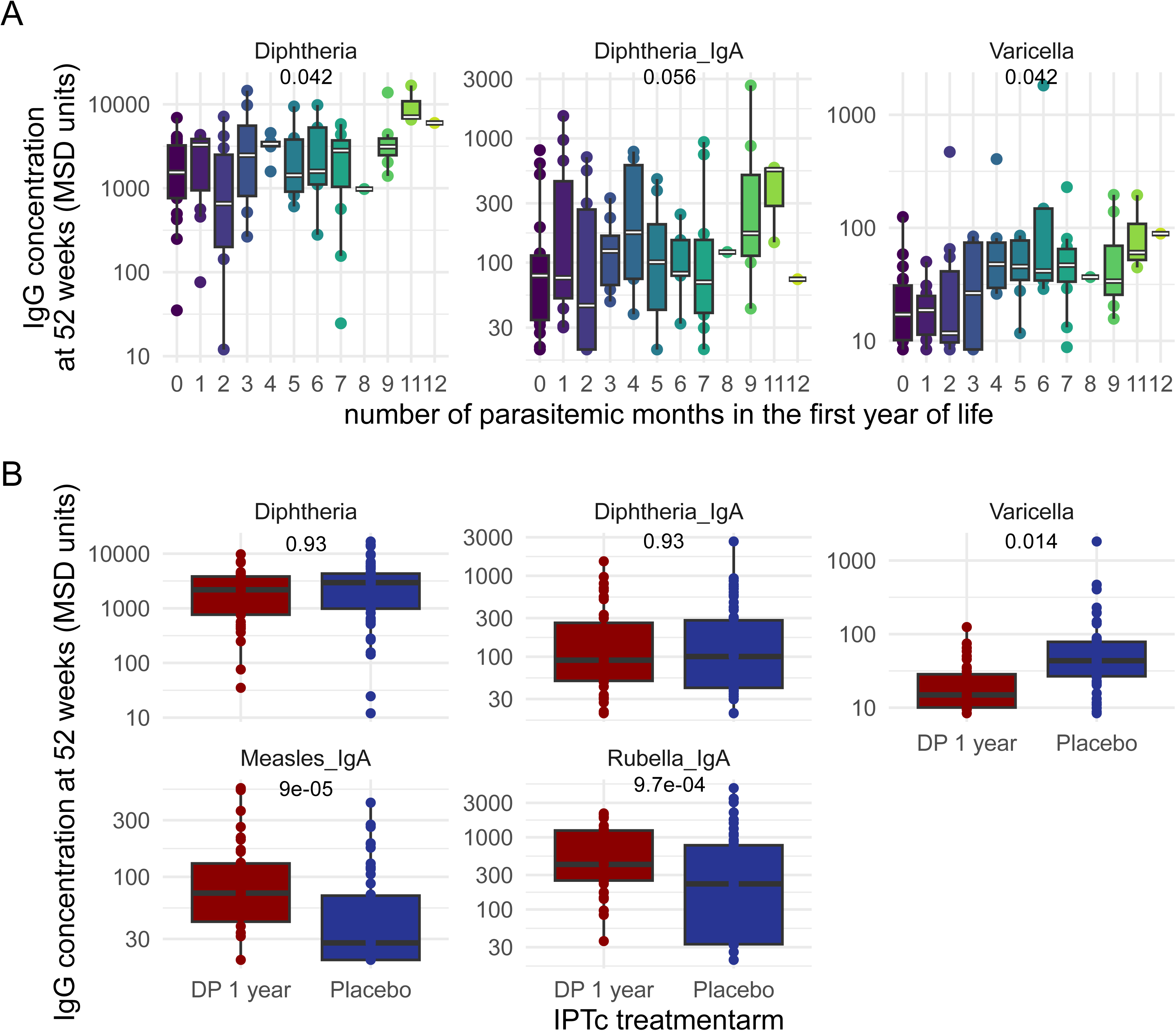
Cumulative *Plasmodium* exposure and chemoprevention treatment arm are associated with diphtheria and varicella antibody responses. (A) Diphtheria IgG, diphtheria IgA, and varicella IgG concentrations at 52 weeks, stratified by the cumulative number of parasitemic months experienced during the first year of life (0–11 months; color scale). (B) Antibody concentrations to diphtheria (IgG and IgA), varicella, measles IgA, and rubella IgA, compared between infants randomized to one year of dihydroartemisinin-piperaquine (DP) chemoprevention versus placebo.

**Figure 7.**
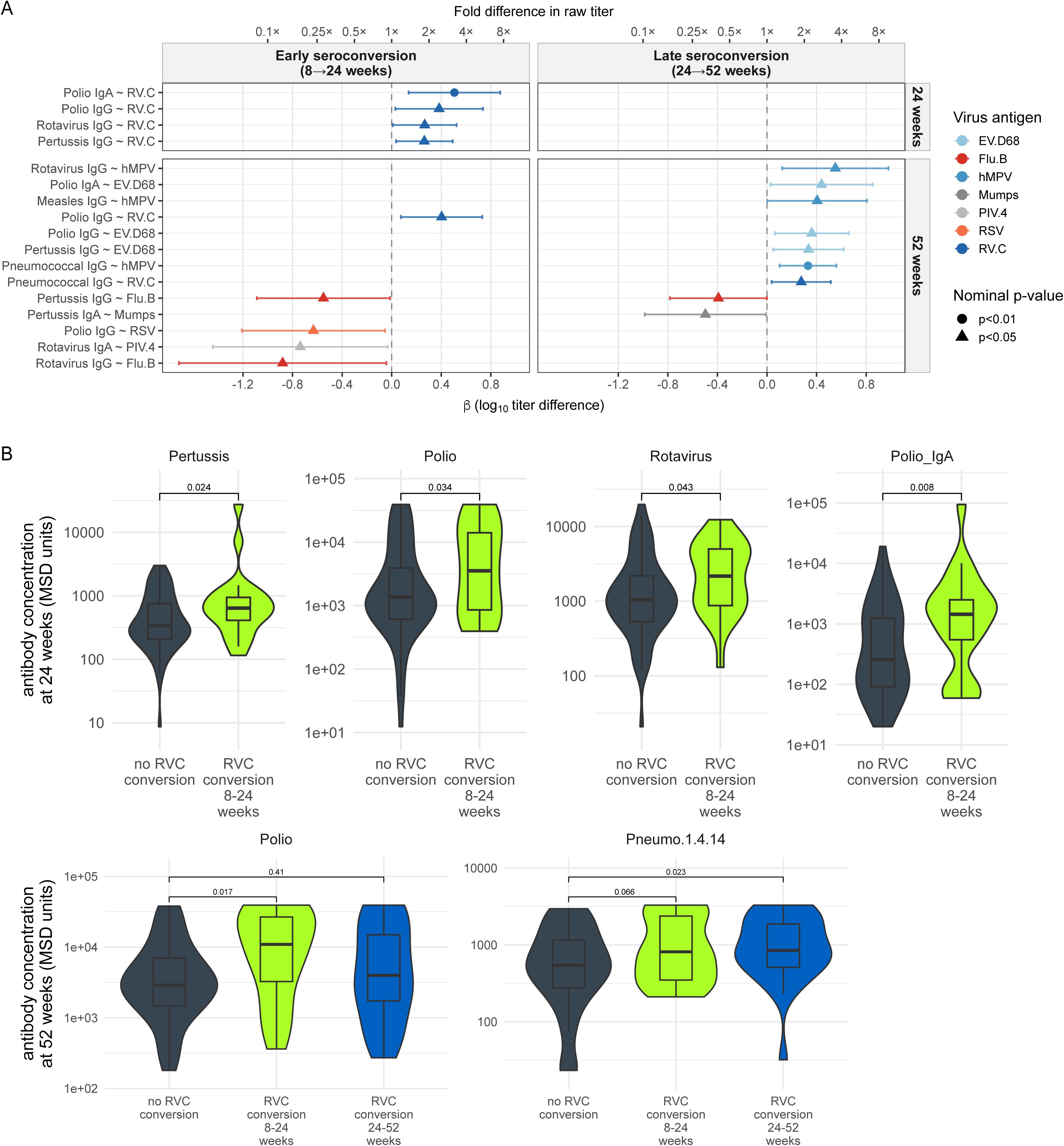
Respiratory viral seroconversion is associated with higher vaccine antibody titers. **(A)** Forest plot showing associations between early (8–24 weeks) and late (24–52 weeks) viral seroconversion and log10-transformed vaccine antibody titers at 24 and 52 weeks of age. Each row represents a single vaccine antigen × virus antigen association estimated by linear regression. Points show β coefficients (log10 titer difference between seroconverters and non-seroconverters) and horizontal bars show 95% confidence intervals. The dashed vertical line indicates no difference (β = 0). Circle and triangle symbols denote nominal p < 0.01 and p < 0.05 respectively. No associations survived FDR correction across the full family of tests. **(B)** Violin plots showing raw antibody concentrations (MSD units) at 24 weeks (upper row) and 52 weeks (lower row) for nominally significant RVC associations, stratified by RVC seroconversion status: no RVC conversion, RVC conversion between 8 and 24 weeks, and RVC conversion between 24 and 52 weeks. Individual data points are overlaid. P-values shown are from the linear models described in panel A.

Since around half of the children in this cohort were exposed to highly effective chemoprevention, we next compared antibody responses between children who received DP or placebo in the first year of life. Overall, we did not see significant differences in serconversion to viral antigens (Fig. S6). Neither diphtheria IgG nor IgA levels were significantly different between chemoprevention arms. In contrast, we found that children who received DP had significantly lower levels of varicella antibodies (Fig. 6B). Consistent with this, children receiving malaria chemoprevention with DP for one or two years had significantly fewer clinical cases of chickenpox than children receiving placebo (incidence rate ratio = 0.46 (95% CI 0.24–0.87), p = 0.018 for DP vs. no DP). While these differences may be due to differences in exposure, we also found differences in antibody responses to vaccine antigens. We found that children who received DP had higher levels of measles and rubella IgA, whereas IgG levels were similar. While it is unclear if higher IgA levels to these viruses impact protection, this finding could be indicative of an immunomodulatory effect of DP, resulting in differences in antibody class ratios.

In summary, we found that both parasitemia at time of measurement, as well as infection history with *P. falciparum* was associated with higher concentrations of both IgG and IgA recognizing vaccine and viral antigens. This could indicate that *P. falciparum* infection predisposes to respiratory infections; alternatively, it is possible that ongoing immune responses to *P. falciparum* may non-specifically increase antibody responses to other antigens, resulting in higher antibody concentrations to viruses and vaccines.

### Infection history with respiratory viruses is associated with vaccine responses

Next, we examined whether infection history with other pathogens contributed to vaccine antibody responses. We tested associations between early seroconversion (8–24 weeks) to rhinovirus C (RVC) and SARS-CoV-2, the only two viruses with ten or more seroconversions in this window, and vaccine titers at 24 weeks across nine antibodies (excluding measles and rubella, for which the first vaccine dose is administered after 24 weeks). Early RVC seroconversion was positively associated with higher titers across four vaccine antigens: Polio IgA (β = 0.51, 95% CI 0.14–0.88, p = 0.008), Pertussis IgG (β = 0.26, 95% CI 0.03–0.49, p = 0.024), Polio IgG (β = 0.38, 95% CI 0.03–0.74, p = 0.034), and Rotavirus IgG (β = 0.27, 95% CI 0.01–0.52, p = 0.043), corresponding to 1.8–3.2 fold higher titers on the raw scale. No associations survived FDR correction across the 18-test family. However, all four RVC associations were positive in direction, unlikely under a null of random directionality (sign test: 4/4 positive, p = 0.063). No significant associations were observed for SARS-CoV-2.

We repeated this analysis at 52 weeks, including all eleven vaccine antigens and fourteen viral antigens with ten or more seroconversions between 24 and 52 weeks (308 tests). Again, no associations survived FDR correction. The association between early RVC seroconversion and Polio IgG observed at 24 weeks persisted at 52 weeks (β = 0.40, 2.53×, p = 0.017), suggesting a durable lagged effect extending at least six months after initial seroconversion. In addition, late seroconversion to RVC was associated with higher pneumococcal IgG (β = 0.28, p = 0.023). Nominally, late seroconversion to human metapneumovirus (hMPV) and enterovirus D68 (EV.D68), both mucosal respiratory viruses, was associated with higher titers to multiple vaccine antigens: hMPV with Rotavirus IgG (β = 0.55, 3.55×, p = 0.012) and Pneumococcal IgG (β = 0.33, 2.14×, p = 0.005), and EV.D68 with Polio IgG (β = 0.36, 2.30×, p = 0.018) and Pertussis IgG (β = 0.34, 2.16×, p = 0.022). In contrast, early seroconversion to influenza B and RSV was nominally associated with lower vaccine titers at 52 weeks, including Rotavirus IgG (Flu.B: β = −0.88, 0.13×, p = 0.040), Pertussis IgG (Flu.B: β = −0.55, 0.35×, p = 0.045), and Polio IgG (RSV: β = −0.63, 0.23×, p = 0.033). However, with only two to three hits per virus and no sign-test support, these negative associations should be regarded as hypothesis-generating observations rather than a demonstrated pattern.

RVC seroconversion was the most robust and consistent predictor of higher vaccine antibody titers, with directionally uniform associations across six vaccine antigen contrasts spanning both timepoints (sign test: 6/6 positive, p = 0.016).

## Discussion

This longitudinal serological study characterized vaccine antibody acquisition and durability across the first year of life in Ugandan infants, revealing substantial heterogeneity in responses to routine EPI antigens and identifying several host and environmental factors associated with vaccine immunogenicity. Embedding this sub-study within the MICDROP trial allowed us to examine these responses in the context of prospective malaria surveillance, placental histology, and natural exposure to common viruses– a combination of covariates rarely available in vaccine immunogenicity studies. Collectively, our findings identify specific vulnerabilities in the current EPI schedule in this setting, and raise the possibility that early-life infectious exposures, including both *P. falciparum* and common respiratory viruses, may modulate the magnitude of vaccine antibody responses.

Interestingly, we found consistent positive associations between early rhinovirus C seroconversion and antibody responses to multiple vaccine antigens, including polio, pertussis, rotavirus, and pneumococcus, across both 24 and 52 weeks of age. This pattern raises the intriguing possibility that early mucosal viral infections may act as heterologous adjuvants during a critical window of immune development. Nominally, other mucosal and enteric viruses (hMPV, EV.D68) showed similar positive patterns, while systemic respiratory viruses (influenza B, RSV) showed the opposite trend, though with only one to two hits each the latter observations remain hypothesis-generating. Respiratory viral infection is known to induce type I interferon signaling and broad innate immune activation, which could plausibly enhance antigen presentation and B cell help at the time of vaccination, independent of the infecting pathogen itself. The positive association between *P. falciparum* exposure and both vaccine as well as viral antibody responses raises related and distinct questions. Parasite carriage may be associated with an immune state more conducive to viral infection or higher viral loads, with elevated antibody responses reflecting greater cumulative antigen exposure rather than a direct adjuvant effect of malaria itself. Disentangling these possibilities will require studies that characterize concurrent inflammatory and regulatory immune states in the context of longitudinal infection and vaccination histories. We cannot exclude the possibility that shared socioeconomic or household-level factors, such as housing quality and parental health literacy, independently increase exposure to both *P. falciparum* and respiratory viruses, which would produce the observed co-associations without any direct immunological interaction between the two.

We found that only 23.6% of infants retained highly protective diphtheria antibody levels ≥ 0.1 IU / mL, with 86.5% of infants ≥ 0.01 IU / mL, the minimally protective threshold, at one year of age. This is consistent with similar findings in other African pediatric cohorts: Tohme *et al.* found only 35% and 77.2% of children under four years old reached these levels in a large Nigerian study^17^. This critical gap in vaccine-induced immunity is concerning, especially since diphtheria is resurgent in several African countries^18^. The stark contrast with the robust tetanus responses from the same combination vaccine suggests antigen-specific differences in immunogenicity rather than a generalized vaccine failure. Current vaccination schedules may require optimization through additional booster doses or alternative formulations to achieve durable protection in this population^13^.

Pertussis responses in our cohort merit particular attention. Previous work has shown that young children exhibit significantly faster rates of waning IgG to several *B. pertussis* antigens^19^. While no definitive correlate of protection has been described for pertussis vaccination, both IgA^20^ and IgG^21,22^ appear to play important roles. In animal models, IgG recognizing B. pertussis filamentous haemagglutinin has been shown to transudate into the respiratory tract and reduce colonization in the lung and trachea^23^. As IgG levels peaked at 8 weeks and failed to be boosted following immunization in our cohort, childhood vaccination may confer incomplete protection against pertussis in this population. Clinically, pertussis remains an important and prevalent childhood illness in Uganda, despite high vaccination rates^24^, consistent with relatively reduced immunogenicity of this vaccine component. The continued increase in pertussis- and rotavirus-specific IgA from 8 to 24 to 52 weeks is most plausibly explained by natural exposure rather than by maturation of the vaccine-induced response. For rotavirus, this interpretation is well supported by studies conducted in comparable high-transmission settings: placebo-arm data from rotavirus vaccine trials show substantially higher background IgA seropositivity in African infants than in higher-income settings, indicating that natural rotavirus infection occurs early and commonly in these populations, regardless of vaccination status^25^. Breakthrough rotavirus infection in vaccinated infants is similarly well documented in sub-Saharan Africa^26^, consistent with the IgA rise we observed continuing well beyond the primary vaccination series. For pertussis, Uganda’s EPI schedule uses a whole-cell pertussis (wP)-containing formulation (DPT-HepB-Hib), which, unlike acellular vaccines, may itself prime mucosal IgA^27^. We are therefore unable to fully exclude a vaccine-induced contribution to the pertussis IgA rise in this cohort. Nonetheless, given the continued clinical burden of pertussis in Uganda despite high vaccination coverage, and the well-established high transmission intensity of *B. pertussis* in this setting, natural exposure remains a plausible and likely contributor to the IgA kinetics observed here.

Measles and Rubella vaccination also appeared to underperform in this population: in the United States, a single dose is 93% and 97% effective against Measles and Rubella, respectively, whereas only ∼76% and ∼ 74% of Ugandan infants reached protective thresholds against Measles and Rubella, respectively. We did not systematically assess nutritional status, including vitamin A sufficiency and stunting, which are among the best-established determinants of measles vaccine immunogenicity in sub-Saharan African children. Malnutrition can impair both primary vaccine responses and antibody durability, and may indeed explain reduced immunogenicity in this setting.

Viral seroconversion patterns offered additional insight into local transmission dynamics and their interaction with early-life exposures. By one year of age, 10/89 (11.2%) of infants had seroconverted to mumps. This high incidence of mumps seroconversion in such young children implies high community transmission intensity and may consideration of mumps vaccination in future EPI schedule revisions. We similarly observed 12/89 (13.5%) serconversion to varicella. While this study was not powered to test associations between treatment arm and seroconversion, we did find that DP was associated with lower concentrations of antibody. Additionally, we found that children who received DP during infancy were significantly less likely to present with symptomatic chicken pox during clinic visits. This suggests that *P. falciparum* exposure in early life may predispose infants to varicella infection, leading to higher rates of both seroconversion and clinical illness, consistent with the broader pattern described above, in which *P. falciparum* exposure appears to shape susceptibility to co-occurring infections.

Curiously, we found significant correlations with maternal gravidity, with children born to primigravid mothers had significantly lower antibody concentrations to measles, rotavirus, tetanus, varicella and parainfluenza virus 2 (PIV2) at 8 weeks of age compared to those born to *multigravidae*. Very few studies have directly tested the impact of gravidity on antibody transfer, outside of contexts of placental pathology, generally finding a lack of association between parity / gravidity and antibody transfer in cord blood^28–30^. In Uganda, primigravidity is associated with a higher incidence of adverse birth outcomes^31,32^ and we have shown recently that low birth weight and prematurity are associated with lower IgG responses to PCV10 vaccination^33^. Nonetheless, neither gestational age, maternal age at delivery, nor placental malaria fully explained the association between lower antibody levels at 8 weeks and primigravid mothers presented here. It is possible that our results are driven by epidemiological factors: for example, the number of children already present in the household during pregnancy, and their associated microbes, may impact both maternal antibody responses that are transferred, as well as post-birth antibody responses in the infant themselves.

## Strengths and limitations

This study benefits from several important design features. The longitudinal structure, with serial antibody measurements from 8 weeks through one year of age, allowed us to characterize both the acquisition and durability of vaccine responses within the same infants, rather than relying on cross-sectional designs. Embedding this serological sub-study within the MICDROP trial provided unusually rich covariate data, including prospective parasitological surveillance, placental malaria histology, and detailed clinical follow-up, that are rarely available alongside vaccine immunogenicity outcomes. The breadth of antigens measured, spanning EPI vaccine targets, viral seroconversion, and both IgG and IgA isotypes, enabled us contextualize vaccine responses within the broader infectious exposures of early life. Several limitations should be considered when interpreting our findings. First, the study was nested within a trial designed primarily to evaluate malaria chemoprevention, and the serological sub-study was not powered to detect all associations of interest; our findings should be regarded as exploratory and hypothesis-generating. Second, while we identified significant associations between *P. falciparum* exposure and vaccine responses, the cross-sectional nature of some comparisons and the potential for residual confounding preclude causal inference. The mechanisms underlying these associations, whether driven by direct immunomodulatory effects of malaria, co-occurring viral infections, or shared determinants of infection susceptibility, remain to be established. Lastly, total IgG or IgA concentration does not reflect functional antibody quality; viral neutralization capability is often more informative than binding alone. In addition, IgG subclass distribution and post-translational are relevant for long-term protection but were beyond the scope of this study.

Taken together, our findings suggest that vaccine immunogenicity in this Ugandan infant cohort is shaped by the broader infectious and host environment in which vaccination occurs, including concurrent viral infection, *P. falciparum* exposure, and maternal gravidity. The unexpected positive association between heterologous viral infection and vaccine antibody responses, if confirmed in larger and more mechanistically focused studies, would suggest that the timing and infectious context of vaccination may matter as much as the vaccine formulation itself in high-transmission settings. More immediately, our data on antigen-specific durability, particularly the rapid waning of diphtheria antibody and the reduced immunogenicity of pertussis, add to a body of evidence supporting the reconsideration of booster schedules for these antigens in similar populations. More broadly, these findings are a reminder that improving global vaccine effectiveness will require development and testing in the diverse populations where current formulations underperform. Future work directly measuring immune states and functional antibody quality will be needed to establish whether the associations we describe reflect causal, exploitable mechanisms or shared upstream determinants of infection and immune competence.

## Acknowledgements & Funding

The MICDROP clinical trial is funded by the National Institute of Allergy and Infectious Diseases, National Institutes of Health (U01AI155325). Children recruited to MICDROP were born to mothers enrolled in the DPSP study (U01AI141308). PJ is a Woods Family Faculty Scholar in Pediatric Infectious Diseases, FAB was supported by the Walter V. and Idun Berry Postdoctoral Fellowship.

## Materials and Methods

### Cohort Data

Longitudinal plasma samples were collected from 89 Ugandan infants at three timepoints: 8, 24, and 52 weeks of age, nested within the MICDROP clinical trial (NCT04978272. Children were vaccinated at routine visits scheduled in concordance with the Ugandan Ministry of Health’s recommendation for childhood vaccination. The schedule is shown in supplementary table 1.

### Antibody Assays

Antibodies against a set of common childhood viral infections and a set of viral and bacterial childhood vaccine targets were measured in a multiplexed electrochemiluminescence (ECL) indirect serology format using three kits from MESO SCALE DISCOVERY (MSD): V-PLEX® Respiratory Panel 5, V-PLEX Respiratory Panel 6 and V-PLEX Vaccine Panel 1. The kits employ MSD 10-Spot 96-Well MULTI-ARRAY® plates with an array of up to 10 immobilized target antigens in each well. The arrays are printed on integrated carbon electrodes within each well that serve as the solid-phase support for the arrays, and the source of electrical energy for carrying out ECL measurements. Table 1 lists the viral and bacterial targets of each panel. Supplementary Table 2 provides details on the specific antigen construct used for each target. Testing with the V-PLEX Vaccine Panel 1 used a pre-production lot that is identical to the final product except that it included an additional array element for measuring responses to pneumococcal vaccines.

Assays were performed according to the protocol in the package inserts for the kits, which is briefly summarized as follows: The assay plates were blocked by adding MSD Blocker A and incubating for 1 hour. The plates were washed and 25 µL of diluted sample was added to each well and incubated with shaking for 2 hours. The plates were washed and 25 µL of a labeled (SULFO-TAG™ ECL label) anti-human IgG (MSD D21ADF) or anti-human IgA (MSD D21ADE) detection antibody was added to each well, and the plates were incubated with shaking for 1 hour. The plates were washed and 1X MSD GOLD™ Read buffer B was added. Plates were then analyzed using an MSD MESO® SECTOR S 600MM ECL plate reader to generate ECL signals for each array element.

Samples were diluted 1:5,000 in MSD Diluent 100 prior to measurement with Respiratory Panels 5 and 6. For all targets except *S. pneumococcus*, samples were diluted 1:1,000 in MSD Diluent 100 prior to measurement with Vaccine Panel 1. The reported antibody levels to the pneumococcal capsular polysaccharide antigen came from a repeat set of runs of Vaccine Panel 1 that used samples diluted in MSD Diluent 100 plus 10 ug/mL pneumococcal cell wall polysaccharide (CWPS, SSI Diagnostica) to prevent interference from anti-CWPS antibodies. IgG antibodies were measured for all panels/assays. IgA antibodies were only measured for Vaccine Panel 1 for samples without CWPS blocker.

### Antibody Quantitation

Antibodies were quantified, following the protocols in the kit product inserts, against calibration standards provided with the respective kits having assigned values for each pathogen target and Ig type in MSD Arbitrary Units per mL (AU/mL). Serial dilution of the calibration standards was used to generate 7-point calibration curves that were run on each assay plate. ECL signals from the calibrator dilution series were fit to a four-parameter-logistic (4PL) model using 1/Y^2^ weighting. Samples that quantified below the Lower Limit of Quantitation (LLOQ) or Upper Limit of Quantitation (ULOQ), as defined by the product inserts, were set to the LLOQ or ULOQ value, respectively, for plotting and statistical analysis. Reported concentrations were further corrected for sample dilution.

### Estimation of Protective IgG Thresholds

Protective IgG thresholds in AU/mL were estimated for six of the vaccine antigens in Vaccine Panel 1 that had (i) WHO International Standards comprising pooled serum samples with assigned IgG concentrations in International Units per mL (IU/mL) and (ii) literature reports with predicted protective thresholds for IgG in IU/mL.^9,10^ The WHO International Standards were obtained from the UK National Institute for Biological Standards and Controls (NIBSC). Each of these standards was tested with Vaccine Panel 1 at three dilutions within the linear range of the respective assays. The measured concentration in MSD AU/mL at the three dilutions was averaged and used to calculate a multiplicative conversion factor to convert IU to AU. Supplementary Table 3 lists the six vaccine antigens, the WHO International Standards, the conversion factors and estimates of protective IgG levels calculated from literature values using the conversion factors.

### Statistical Analysis

All antibody concentrations were log□□-transformed prior to analysis. Parasitological data, including concurrent qPCR-quantified parasite density (log□□-transformed with a pseudocount of 0.001) and longitudinal infection history, were integrated from the parent trial database. Maternal metadata including treatment arm, gravidity, age, and placental histopathology were linked by subject ID.

### Covariate screening

Prior to primary analyses, we systematically evaluated whether infant sex, household wealth category, and maternal education level were associated with longitudinal antibody trajectories. For each antigen, we fitted linear mixed-effects models using lmerTest::lmer with log□□-transformed AU as the outcome, a fixed effect of timepoint, and a random intercept for subject to account for repeated measures. Each covariate was tested by comparing a base model (timepoint only) to additive and interaction models using likelihood ratio tests (LRT) via the *anova* function, with Benjamini-Hochberg (BH) false discovery rate (FDR) correction applied across antigens. None of sex, wealth, or education significantly improved model fit or yielded FDR-corrected associations with antibody levels, and these covariates were excluded from subsequent primary models.

### Testing interactions with infant and maternal characteristics

The effect of infant treatment arm (monthly DP, or placebo) on longitudinal antibody trajectories was assessed using linear mixed-effects models with a treatment arm × timepoint interaction and a random intercept for subject. Estimated marginal means were computed using the emmeans package and pairwise contrasts were extracted at each timepoint. FDR correction was applied within each timepoint contrast group. Similarly, maternal gravidity was examined using an analogous approach, by fitting mixed-effects models with a gravidity × timepoint interaction and a random intercept for subject. To determine whether the association with gravidity was explained by placental malaria, gestational age, or maternal age, we refitted models including those covariates.

### Seroconversion and viral infection interactions

Seroconversion was defined for viral antigens using a combination of fold-change thresholds: a log₂ fold-change greater than 1 (or a standardized fold-change z-score >2) between 8 and 24 weeks was classified as early seroconversion; a log₂ fold-change greater than 2 between 24 and 52 weeks was classified as late seroconversion. Infants not meeting either criterion were classified as non-converters. To evaluate whether early viral seroconversion was associated with vaccine antibody responses, we fit separate linear regression models for each vaccine antigen × virus antigen combination, stratified by timepoint. Seroconversion status was defined on a per-timepoint basis: for the 24-week analysis, children were classified as early seroconverters if their fold-change in virus-specific antibody between 8 and 24 weeks met the pre-specified seroconversion threshold, and as non-seroconverters otherwise, such that children who seroconverted later (24–52 weeks) were correctly included in the non-seroconverter reference group at 24 weeks. For the 52-week analysis, both early (8–24 week) and late (24–52 week) seroconversion contrasts were included, with non-seroconverters as the reference in both cases. Vaccine titer (log10-transformed) was modeled as a function of seroconversion status. Models were fit using ordinary least squares; each stratum contained at most one observation per child, so no random effects for child identity were required. To ensure adequate power in each stratum, models were only fit for virus antigens with at least ten seroconverters in the relevant window, yielding two virus antigens at 24 weeks (rhinovirus C and SARS-CoV-2; 18 total tests) and fourteen at 52 weeks (308 total tests). P-values from all models within each timepoint were pooled and corrected for multiple comparisons using the Benjamini-Hochberg false discovery rate procedure, applied separately within each timepoint family. Given the modest number of tests at 24 weeks, directional consistency of associations was additionally assessed using an exact binomial sign test. All analyses were performed in R using the tidyverse and broom packages.

---

**Figure S1.**
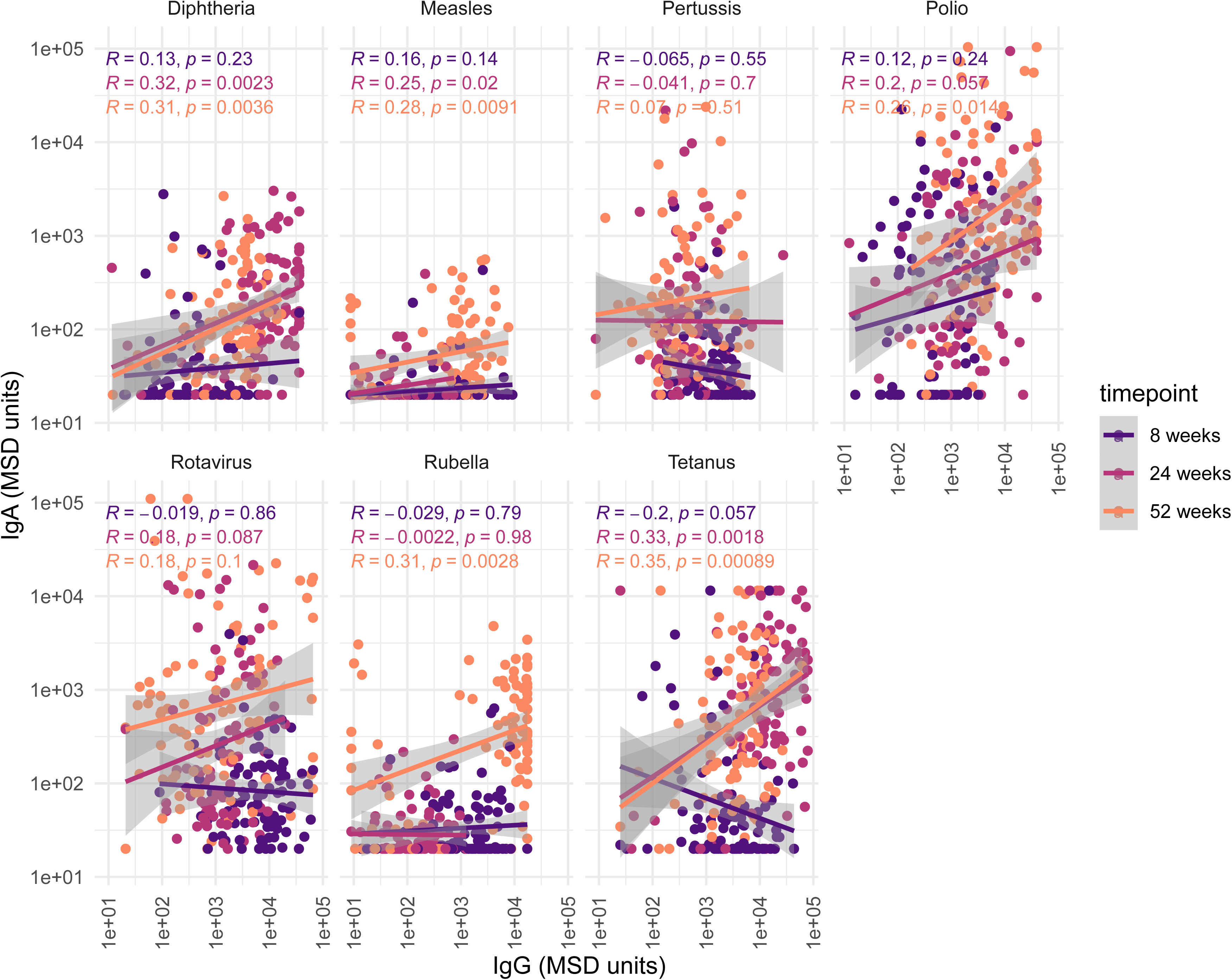
Correlation between IgA and IgG responses across vaccine antigens. Scatterplots of IgA versus IgG concentration (MSD units, log10 scale) for diphtheria, measles, pertussis, polio, rotavirus, rubella, and tetanus, colored by timepoint (8, 24, 52 weeks). R and p-values indicate spearman correlation coefficients across all timepoints combined. Lines show linear regression fit with 95% confidence intervals.

**Figure S2.**
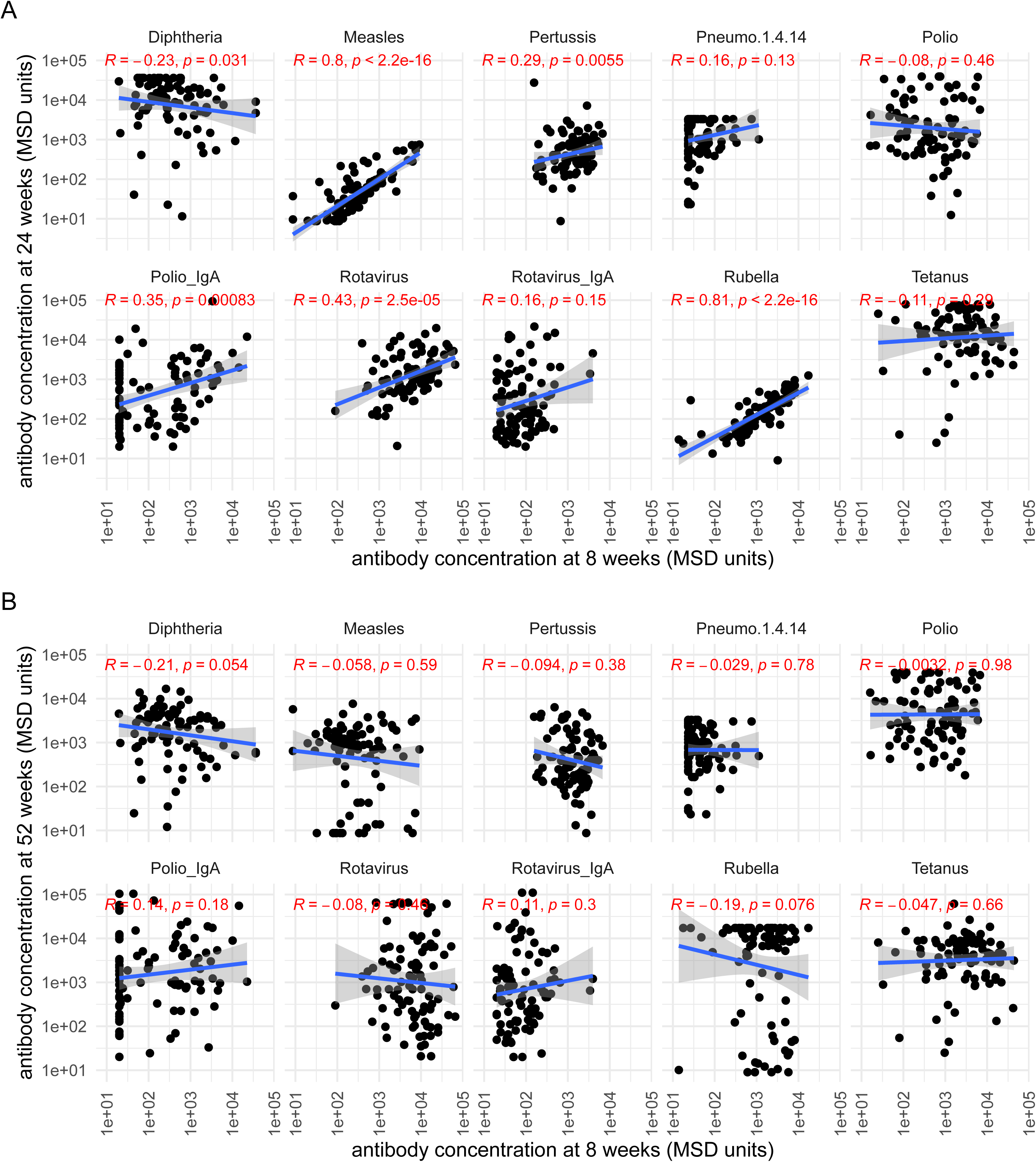
Antibody concentrations at 8 weeks correlate with 24-week but not 52-week titers. Scatterplots of antibody concentration at 8 weeks versus 24 weeks (top) and versus 52 weeks (bottom) for diphtheria, measles, pertussis, pneumococcal serotype 1.4.14, polio (IgG and IgA), rotavirus (IgG and IgA), rubella, and tetanus. R and p-values indicate Spearman correlation coefficients.

**Figure S3.**
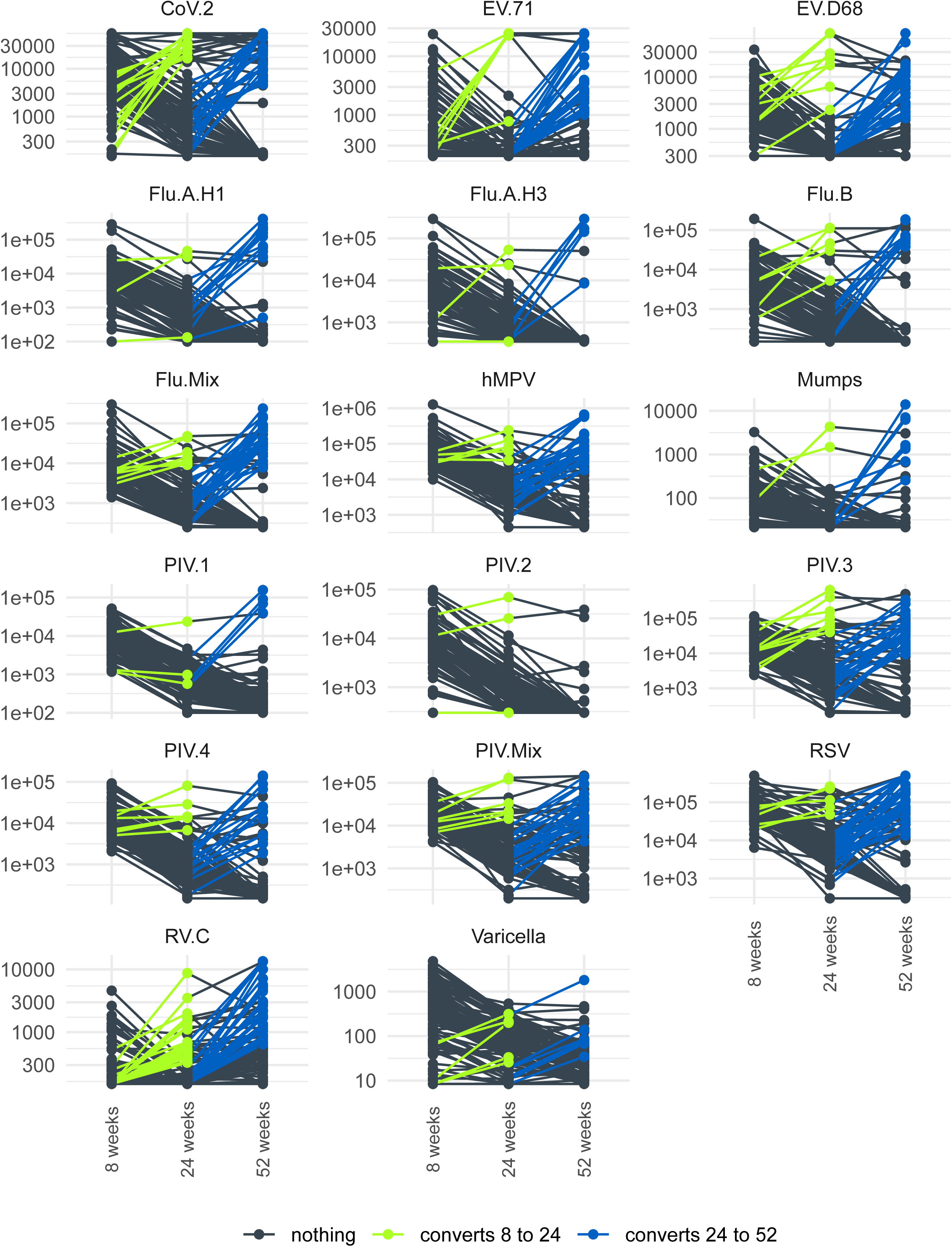
Individual antibody trajectories for all viral antigens assessed. Full longitudinal seroconversion trajectories from 8 to 52 weeks of age for all respiratory and other viral antigens measured, including SARS-CoV-2, enteroviruses (EV.71, EV.D68), influenza strains (Flu.A.H1, Flu.A.H3, Flu.B, Flu.Mix), hMPV, mumps, parainfluenza viruses (PIV.1–4, PIV.Mix), RSV, rhinovirus C, and varicella. Lines are colored by seroconversion status: dark grey, no seroconversion; green, seroconversion between 8 and 24 weeks; blue, seroconversion between 24 and 52 weeks.

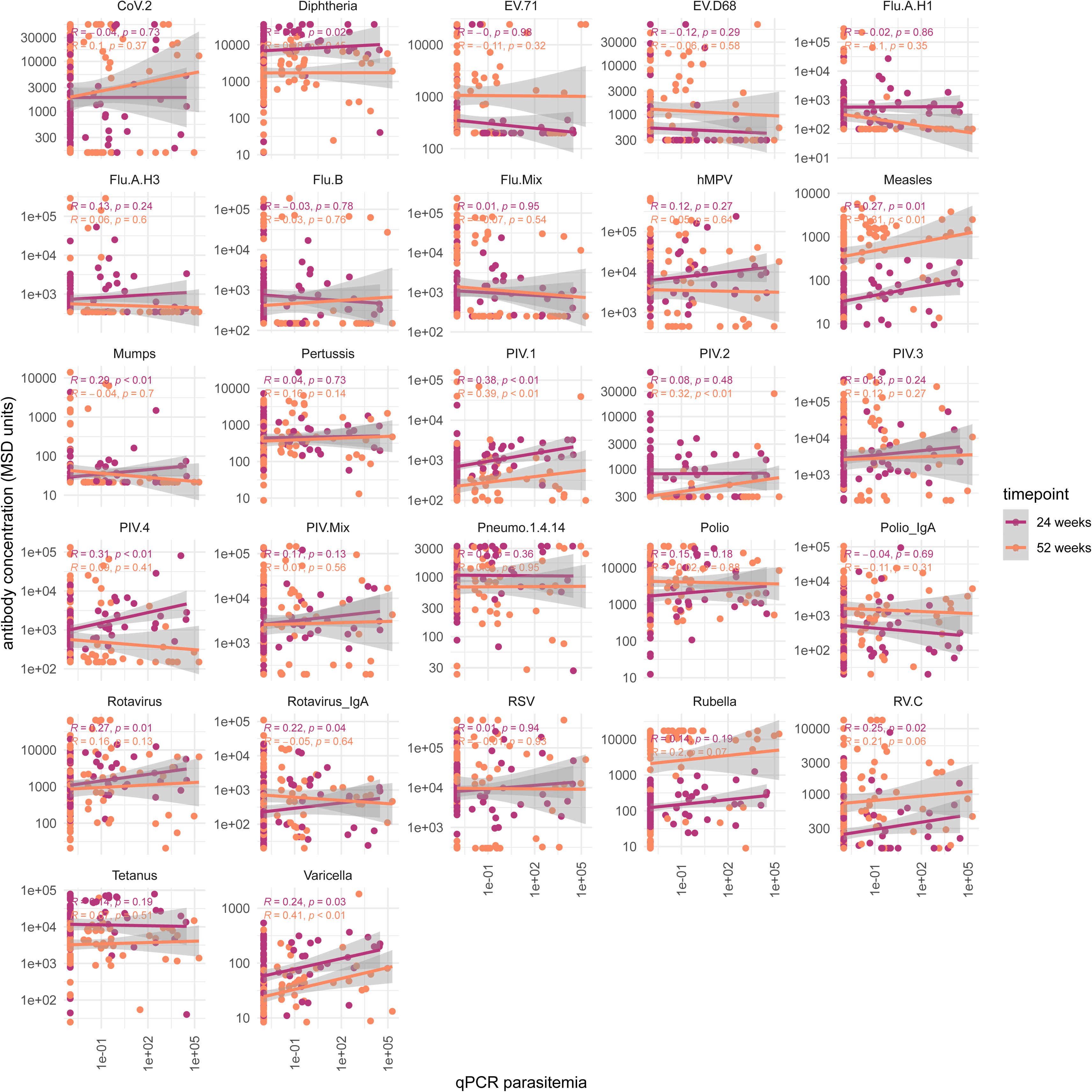

**Figure S5.**
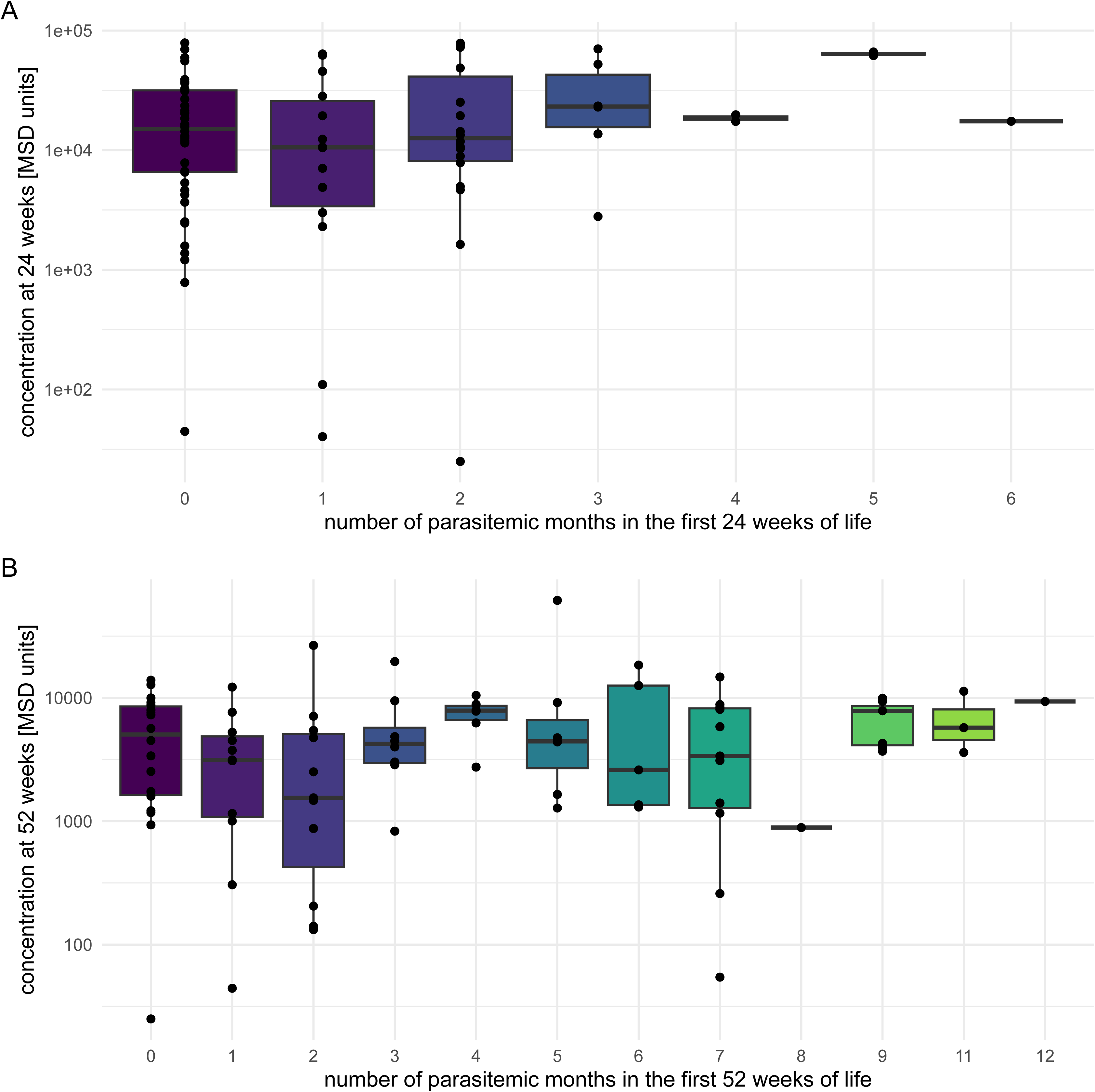
Viral seroconversion does not differ significantly by malaria chemoprevention treatment arm. Cumulative seroconversion by in first year of for all viral antigens assessed, stratified by randomization to dihydroartemisinin-piperaquine (DP, one year) versus placebo. No significant differences were observed between treatment arms for any antigen (proportion test with Yates’ continuity correction); RV.C approached but did not reach significance (p ≈ 0.11).

**Figure S6.**
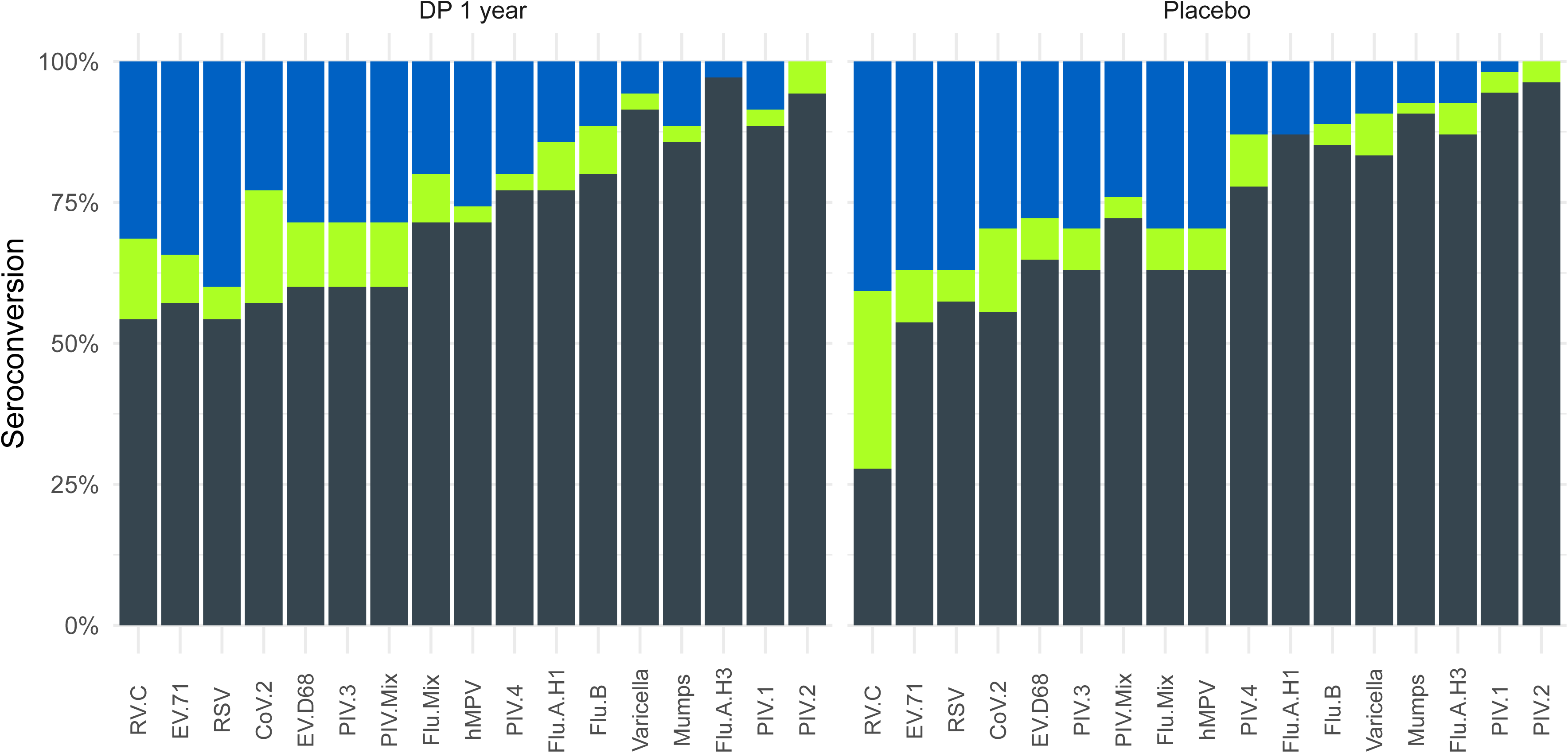
Cumulative *Plasmodium* exposure is not associated with tetanus IgG concentrations. Tetanus IgG concentration (MSD units, log10 scale) at 24 weeks (top) and 52 weeks (bottom) of age, stratified by the cumulative number of parasitemic months experienced up to each timepoint.

**Supplementary Table 1.**
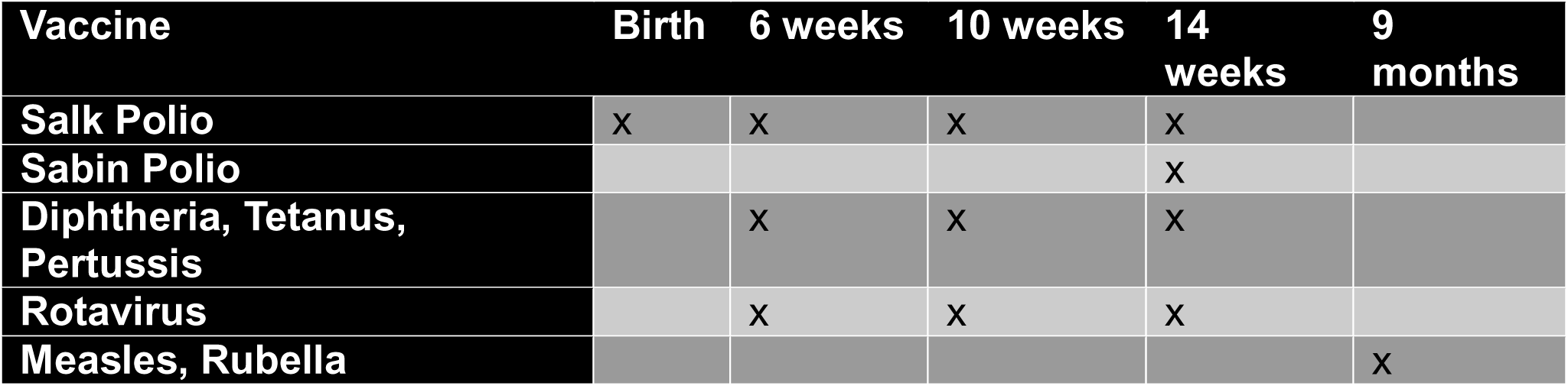
Ugandan vaccine schedule in the first year of life for the vaccines assayed.

**Supplementary Table 2.** Detailed description of the viral and bacterial antigens incorporated into each assay spot in Respiratory Panels 5 and 6, and Vaccine Panel 1.

| Antigens | Antigen Description | Panel Name |
| --- | --- | --- |
| **S. Pneumoniae** | Mix of capsular polysaccharides from serotypes 1, 4 and 14 | Vaccine Panel 1 |
| **Measles HA** | Measles Hemagglutinin | Vaccine Panel 1 |
| **Polio 3 VP1** | Viral protein 1 of the Type 3 poliovirus capsid | Vaccine Panel 1 |
| **mumps HN** | mumps virus Hemagglutinin-Neuraminidase | Vaccine Panel 1 |
| **VZV-gE** | Varicella-zoster virus glycoprotein E | Vaccine Panel 1 |
| **Rubella E1** | Rubella E1 glycoprotein | Vaccine Panel 1 |
| **Diphtheria Toxoid \*** | Toxoid of Diphtheria Toxin produced by *Corynebacterium diphtheria* | Vaccine Panel 1 |
| **Tetanus Toxoid \*** | Toxoid of Tetanus Toxin produced by *Clostridium tetani* | Vaccine Panel 1 |
| **Rotavirus G4 VP6** | VP6 capsid protein from the Rotavirus G4 genotype | Vaccine Panel 1 |
| **B. pertussis FHA 30** | Filamentous Hemagglutinin from *Bordetella pertussis* | Vaccine Panel 1 |
| **SARS-CoV-2 Spike (BA.5)** | Severe Acute Respiratory Syndrome Coronavirus 2 Spike Protein Omicron variant BA.5 | Respiratory Panel 5 |
| **EV-A71 VLP** | Enterovirus 71 virus-like particle | Respiratory Panel 5 |
| **EV-D68 VLP** | Enterovirus 68 virus-like particle | Respiratory Panel 5 |

|  |  |  |
| --- | --- | --- |
| **Flu A/B (mix) HA** | Mix of Flu A/Wisconsin/2019 H1, Flu A/Darwin/2021 H3, and Flu B/Austria/2021 HA | Respiratory Panel 5 |
| **Flu A/Darwin/2021 H3** | Influenza A Hemagglutinin Protein from A/Darwin/6/2021 (H3N2) | Respiratory Panel 6 |
| **Flu A/Wisconsin/2019 H1** | Influenza A Hemagglutinin Protein from A/Wisconsin/588/2019 (H1N1) | Respiratory Panel 6 |
| **Flu B / Austria / 2021 HA** | Influenza B Hemagglutinin Protein from B/ Austria/1359417/2021 (B Victoria L) | Respiratory Panel 6 |
| **hMPV F** | Metapneumovirus F protein | Respiratory Panel 5 |
| **hPIV1 F** | Parainfluenza virus type 1 F protein | Respiratory Panel 6 |
| **hPIV2 F** | Parainfluenza virus type 1 F protein | Respiratory Panel 6 |
| **hPIV13F** | Parainfluenza virus type 1 F protein | Respiratory Panel 6 |
| **hPIV4 F** | Parainfluenza virus type 1 F protein | Respiratory Panel 6 |
| **hPIV(1-4)** | Mix of Parainfluenza virus type 1, 2, 3, and 4 F protein | Respiratory Panel 5 |
| **hRV-C VP0** | Rhinovirus C VPO | Respiratory Panel 5 |
| **RSV Pre-Fusion F** | Respiratory Syncytial Virus Pre-Fusion F Protein | Respiratory Panel 5 |

**Supplementary Table 3.**
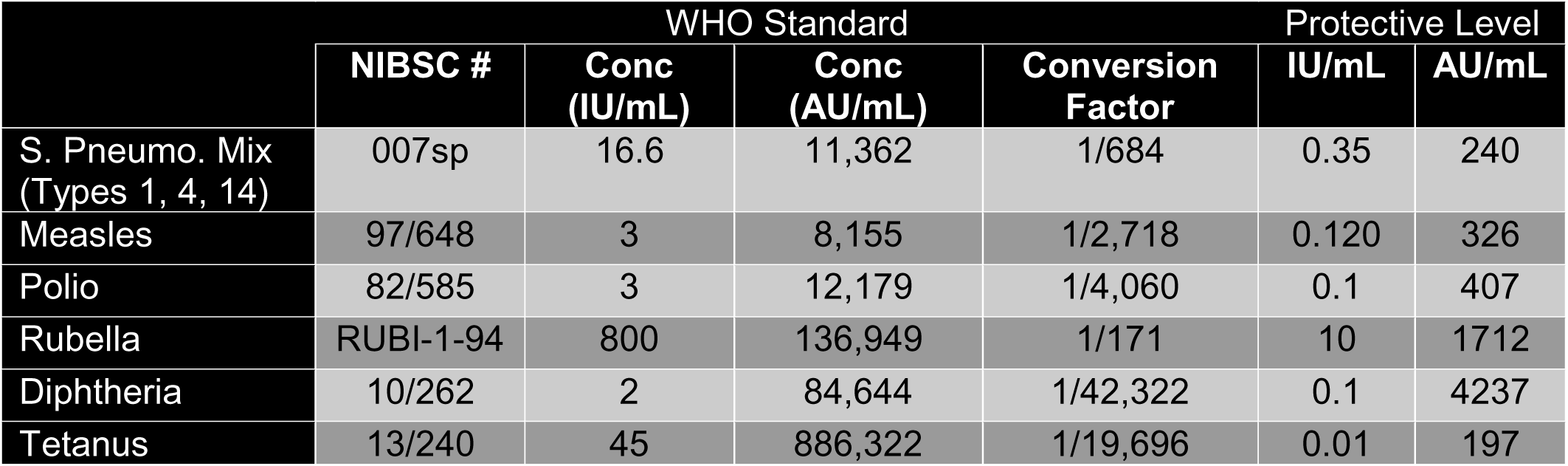
List of WHO International Standards used to convert MSD arbitrary units (AU) to international units (IU). The table lists the NIBSC catalog number of each standard, the assigned concentration in IU/mL, the measured concentration using Vaccine Panel 1 in AU/mL and the multiplicative conversion factor for converting AU to IU. The table also provides a literature estimate of the IgG level (in IU/mL) needed for protection, and the corresponding value in AU/mL calculated using the IU to AU conversion factor. The listed concentration of the pneumo standard in IU/mL is the average of the WHO assigned concentrations for types 1, 4 and 14, where 1 IU equals 1 µg of specific IgG per mL. The listed concentration of the polio standard is the assigned concentration for type 3 polio.

|  | WHO Standard |  |  |  | Protective Level |  |
| --- | --- | --- | --- | --- | --- | --- |
|  | NIBSC # | Conc (IU/mL) | Conc (AU/mL) | Conversion Factor | IU/mL | AU/mL |
| S. Pneumo. Mix (Types 1, 4, 14) | 007sp | 16.6 | 11,362 | 1/684 | 0.35 | 240 |
| Measles | 97/648 | 3 | 8,155 | 1/2,718 | 0.120 | 326 |
| Polio | 82/585 | 3 | 12,179 | 1/4,060 | 0.1 | 407 |
| Rubella | RUBI-1-94 | 800 | 136,949 | 1/171 | 10 | 1712 |
| Diphtheria | 10/262 | 2 | 84,644 | 1/42,322 | 0.1 | 4237 |
| Tetanus | 13/240 | 45 | 886,322 | 1/19,696 | 0.01 | 197 |

## References

1. van Dorst, M. M. A. R. et al. Immunological factors linked to geographical variation in vaccine responses. Nat Rev Immunol 1–14 (2023) doi:10.1038/s41577-023-00941-2.

2. Cunnington, A. J. & Riley, E. M. Suppression of vaccine responses by malaria: Insignificant or overlooked? Expert Review of Vaccines 9, 409–429 (2010).

3. Zirimenya, L. et al. The Effect of Malaria on Responses to Unrelated Vaccines in Animals and Humans: A Systematic Review and Meta-Analysis. Parasite Immunol 46, e13067 (2024).

4. Sbarra, A. N. et al. Assessing the Effect of Malaria Exposure History on Tetanus Antibody Waning Rates Among Children in Jinja and Tororo Districts, Uganda. J Infect Dis jiag255 (2026) doi:10.1093/infdis/jiag255.

5. Nickbakhsh, S. et al. Virus–virus interactions impact the population dynamics of influenza and the common cold. Proceedings of the National Academy of Sciences 116, 27142–27150 (2019).

6. Wu, A., Mihaylova, V. T., Landry, M. L. & Foxman, E. F. Interference between rhinovirus and influenza A virus: a clinical data analysis and experimental infection study. The Lancet Microbe 1, e254–e262 (2020).

7. McFarlane, A. J. et al. Enteric helminth-induced type-I interferon signalling protects against pulmonary virus infection through interaction with microbiota. J Allergy Clin Immunol 140, 1068–1078.e6 (2017).

8. Tapela, K. et al. Cellular immune response to SARS-CoV-2 and clinical presentation in individuals exposed to endemic malaria. Cell Rep 43, 114533 (2024).

9. Zhang, H. et al. Mucosal vaccination in mice provides protection from diverse respiratory threats. Science 0, eaea1260 (2026).

10. Kollmann, T. R. & Marchant, A. Towards Predicting Protective Vaccine Responses in the Very Young. Trends Immunol 37, 523–534 (2016).

11. Plotkin, S. A. Correlates of Protection Induced by Vaccination. Clin Vaccine Immunol 17, 1055–1065 (2010).

12. Diedrich, S., Claus, H. & Schreier, E. Immunity status against poliomyelitis in Germany: Determination of cut-off values in International Units. BMC Infect Dis 2, 2 (2002).

13. Diphtheria vaccines: WHO position paper – August 2017. https://www.who.int/publications/i/item/who-wer9231.

14. Danilova, E., Jenum, P. A., Skogen, V., Pilnikov, V. F. & Sjursen, H. Antidiphtheria Antibody Responses in Patients and Carriers of Corynebacterium diphtheriae in the Arkhangelsk Region of Russia. Clin Vaccine Immunol 13, 627–632 (2006).

15. CDC. Measles Vaccination. Measles (Rubeola) https://www.cdc.gov/measles/vaccines/index.html (2026).

16. Sbarra, A., et al. Assessing the effect of malaria exposure history on tetanus antibody waning rates among children in Jinja and Tororo Districts, Uganda. Preprint at 10.31219/osf.io/72gpq_v1 (2025).

17. Tohme, R. A. et al. Tetanus and Diphtheria Seroprotection among Children Younger Than 15 Years in Nigeria, 2018: Who Are the Unprotected Children? Vaccines (Basel) 11, 663 (2023).

18. Diphtheria - African Region (AFRO). https://www.who.int/emergencies/disease-outbreak-news/item/DON588.

19. van Twillert, I. et al. Impact of age and vaccination history on long-term serological responses after symptomatic B. pertussis infection, a high dimensional data analysis. Sci Rep 7, 40328 (2017).

20. Hellwig, S. M. M., van Spriel, A. B., Schellekens, J. F. P., Mooi, F. R. & van de Winkel, J. G. J. Immunoglobulin A-Mediated Protection against Bordetella pertussis Infection. Infect Immun 69, 4846–4850 (2001).

21. Cherry, J. D., Gornbein, J., Heininger, U. & Stehr, K. A search for serologic correlates of immunity to Bordetella pertussis cough illnesses. Vaccine 16, 1901–1906 (1998).

22. Storsaeter, J., Hallander, H. O., Gustafsson, L. & Olin, P. Levels of anti-pertussis antibodies related to protection after household exposure to Bordetella pertussis. Vaccine 16, 1907–1916 (1998).

23. Kimura, A., Mountzouros, K. T., Relman, D. A., Falkow, S. & Cowell, J. L. Bordetella pertussis filamentous hemagglutinin: evaluation as a protective antigen and colonization factor in a mouse respiratory infection model. Infect Immun 58, 7–16 (1990).

24. Kayina, V. et al. Pertussis Prevalence and Its Determinants among Children with Persistent Cough in Urban Uganda. PLoS One 10, e0123240 (2015).

25. Cunliffe, N. et al. Early exposure of infants to natural rotavirus infection: a review of studies with human rotavirus vaccine RIX4414. BMC Pediatr 14, 295 (2014).

26. Simwaka, J. et al. Rotavirus breakthrough infections responsible for gastroenteritis in vaccinated infants who presented with acute diarrhoea at University Teaching Hospitals, Children’s Hospital in 2016, in Lusaka Zambia. PLOS ONE 16, e0246025 (2021).

27. Hendrikx, L. H. et al. Serum IgA Responses against Pertussis Proteins in Infected and Dutch wP or aP Vaccinated Children: An Additional Role in Pertussis Diagnostics. PLoS One 6, e27681 (2011).

28. Brair, M.-E., Brabin, B., Milligan, P., Maxwell, S. & Hart, C. A. Reduced transfer of tetanus antibodies with placental malaria. The Lancet 343, 208–209 (1994).

29. Okoko, B. J. et al. The Influence of Placental Malaria Infection and Maternal Hypergammaglobulinemia on Transplacental Transfer of Antibodies and IgG Subclasses in a Rural West African Population. J Infect Dis 184, 627–632 (2001).

30. Doroudchi, M., Samsami Dehaghani, A., Emad, K. & Ghaderi, A. Placental transfer of rubella-specific IgG in fullterm and preterm newborns. Int J Gynaecol Obstet 81, 157–162 (2003).

31. Bater, J. et al. Predictors of low birth weight and preterm birth in rural Uganda: Findings from a birth cohort study. PLoS One 15, e0235626 (2020).

32. Kusolo, R. et al. Prevalence, trends, and maternal risk factors of adverse birth outcomes from a hospital-based birth defects surveillance system in Kampala, Uganda, 2015–2022. BMC Pregnancy Childbirth 25, 408 (2025).

33. Musinguzi, K. et al. Low birthweight and prematurity, but not malaria chemoprevention, are associated with reduced pneumococcal vaccine immunogenicity in Ugandan infants. Preprint at 10.64898/2026.05.17.26353405 (2026).

